# Focused Ultrasound Blood-Tumor Barrier Opening Rapidly Augments Intratumor CD4 and CD8 T Cell Representation in a Genetically Engineered Mouse Model of Glioma

**DOI:** 10.64898/2026.07.26.740799

**Authors:** Katherine M. Nowak, Matthew R. Hoch, William B. Gillespie, Claire A. Conarroe, Victoria R. Breza, Tanya Cruz, Eden N. Gordon, Catherine M. Gorick, Tajie H. Harris, Joshua D. Wythe, Richard J. Price

## Abstract

Glioblastoma (GBM) is a highly aggressive primary brain tumor that remains difficult to treat due in part to its disorganized and heterogeneous vasculature, known as the blood-tumor barrier (BTB), which limits therapeutic delivery and beneficial immune cell infiltration. Focused ultrasound (FUS) with microbubbles (MBs) can transiently disrupt the BTB to enhance drug delivery and may induce sterile inflammation (SI) that can beneficially remodel the tumor immune landscape. However, this concept has only been explored in implanted tumor models with modest immune effects. Here, we utilized a physiologically relevant genetically engineered mouse model (GEMM) generated via in utero electroporation targeting *Nf1*, *Tp53*, and *Pten* to study tumor-vascular-immune interactions. This 3x CRISPR-Cas9 GEMM recapitulates key features of human glioma, including infiltrative growth, histopathology, molecular alterations, and stage-dependent blood-brain barrier disruption. FUS+MBs were applied to transiently disrupt the BTB, and MRI confirmed increased vascular permeability in treated tumors. Flow cytometry revealed robust increases in tumor-infiltrating CD4^+^ helper and CD8^+^ effector T cells three days post-FUS treatment, without altering the CD8/Treg ratio. These findings were supported by immunofluorescence imaging. Double-negative and double-positive T cells were detected, but they were not significantly altered by FUS. Ki67 analysis indicated that increased T-cell accumulation was not driven by local proliferation. By seven days post-treatment, immune differences were no longer observed. Collectively, these results demonstrate that FUS-mediated BTB disruption selectively and rapidly enhances lymphocyte infiltration in a clinically relevant glioma model, supporting its potential as a temporally controlled immunomodulatory strategy for GBM.

## Introduction

Glioblastoma (GBM) remains one of the most aggressive and treatment-refractory primary brain malignancies, in part due to the blood-tumor barrier (BTB), which is characterized by abnormal and heterogeneous vasculature[1,2]. The BTB impairs drug delivery and T-cell recruitment and function, restricting therapeutic efficacy and the development of anti-tumor immunity[1,3]. Focused ultrasound (FUS) with microbubbles (MBs) enables noninvasive, site-specific BTB disruption, transiently increasing vascular permeability and facilitating drug access to tumors[4,5]. Beyond enhanced drug delivery, FUS+MBs can trigger a sterile inflammatory response (SIR) marked by glial activation, increased endothelial adhesion molecule expression, and proinflammatory cytokine signaling, collectively priming the tumor microenvironment (TME) for immunomodulation[4,6–13]. However, the immunologic effects of FUS+MB-induced SIR have only been studied in implanted GBM models, which do not fully recapitulate the cellular and molecular heterogeneity, tumor evolution, or immune landscape of human gliomas[14,15]. Findings from syngeneic tumor models have failed to correlate with clinical studies[16,17].

Genetically engineered mouse models (GEMMs) provide a more physiologically relevant system to study tumor-vascular-immune interactions in an immunocompetent context[18,19]. Unlike orthotopic implantation models, GEMMs capture stepwise tumor evolution, infiltrative growth, and immune contexture observed in human GBM[18,20,21]. Endothelial phenotypes in GEMMs better reflect the natural heterogeneity of tumor vasculature, providing insight into mechanisms underlying therapeutic resistance and immune evasion[18,21–23]. By comparison, transplantable brain tumor models often exhibit higher baseline BTB permeability, reflecting fundamental differences in vascular architecture, maturation, and function between these systems[24]. The recently developed in utero electroporation-based 3x CRISPR GEMM targeting *Nf1*, *TP53*, and *Pten* recapitulate key features of human GBM, including diffuse infiltration and progressive blood-brain barrier (BBB) breakdown between postnatal days 65-80 (P65-80)[2].

Here, we report the first application of FUS-mediated BTB opening (BTBO) in a GBM GEMM. We demonstrate that 3x CRISPR gliomas respond to FUS+MBs with increased MRI contrast accumulation, consistent with enhanced vascular permeability. Notably, FUS induced a marked increase in intratumoral T-cell populations three days post-FUS without evidence of increased proliferation. These findings demonstrate that FUS-induced SIR can promote immune infiltration in a clinically relevant glioma model, highlighting the importance of progressive vascular remodeling and BBB dynamics in shaping anti-tumor immune responses.

## Methods

### Genetically Engineered Mouse Model of Glioma

All animal experiments followed University of Virginia guidelines and regulations and were approved by the Animal Care and Use Committee. In utero electroporation was performed at embryonic day 14.5 (E14.5) in CD-1 IGS mice, as previously described[25], with further details provided in *Supplemental Methods* and **Figure S1**. A plasmid mixture was injected into the lateral ventricles of embryos: (1) a CAG-Cas9 vector with sgRNAs targeting murine *Nf1*, *Pten*, and *Tp53*; (2) a GLAST-PiggyBac transposase vector; and (3) a cargo vector with CAG-driven EGFP/luciferase flanked by PiggyBac transposons. For at least 3 days post-surgery, the pregnant dams were administered a daily dose of meloxicam (5 mg/kg). Tumor presence was confirmed on postnatal day 30 (P30) via in vivo bioluminescence imaging after intraperitoneal D-Luciferin (XenoLIght™ D-Luciferin K+ salt) injection (10 μL/gram, 15 mg/mL dissolved in DPBS). Mice were maintained on standard chow and a 12-hour light/12-hour dark cycle until P65-80.

### Focused Ultrasound Application

BBB opening was achieved using a 1.15 MHz FUS Instruments RK-300 system with a single-element transducer (10 ms bursts, 1500 ms burst period, 60 sonications, 1.5-minute sonication duration).

FUS targeting consisted of enough sonication points to allow for full tumor coverage. Control mice that had been electroporated at E14.5, but failed to develop tumors, were treated with an equivalent number of sonication points in the same anatomical location as the tumor-bearing mice.

The FUS Instruments software, operating in the "Blood-brain Barrier" mode, facilitated Passive Cavitation Detection (PCD)-modulated peak negative pressure (PNP). The feedback control system parameters were set as follows: a starting pressure of 0.325 MPa, pressure increment of 0.05 MPa, maximum pressure of 0.45 MPa, 20 sonication baselines without MBs, area under the curve (AUC) bandwidth of 500 Hz, AUC threshold of 8 standard deviations, pressure drop of 0.95, and frequency selection of the subharmonic, first ultraharmonic, and second ultraharmonic. AUC refers to the area under the curve of the frequency spectrum, and the specified bandwidth defines the range around each harmonic frequency over which the spectral power is integrated. Optison^™^ (GE HealthCare) MBs were intravenously injected as a bolus dose (2x10^5^ microbubbles/g body weight). Detailed MRI methods and PCD analyses are provided in *Supplemental Methods*.

### Flow Cytometry

Tissue was processed for flow cytometry as described in *Supplemental Methods*. Single-cell suspensions were transferred to a 96-well U-bottom plate for staining. The samples were centrifuged at 1500 RPM for 5 min and then resuspended in 50 μL of Fc Block, made in FACS buffer (1X PBS, 0.2% BSA, and 2 mM EDTA) with 0.1 μg/ml 2.4G2 Ab (BD Biosciences #553142) and 0.1% rat gamma globulin (Fischer Scientific #PI31885) for 10 min. Fluorescent antibodies were added to each sample to stain for surface markers and Live/Dead™ Blue (1:800, Invitrogen #L23105) for 1 hr at 4°C in the dark. Panel surface markers included: CD45-BUV395 (1:800, BD Biosciences #565967), CD11b-BB700 (1:200, BD Biosciences #566416), MHCII-BUV496 (1:400, BD Biosciences #750281), CD44-BUV737 (1:200, BD Biosciences #612799), CD11c-eFlour™ 506 (1:200, ThermoFisher #69-0114-82), CD3-BUV563 (1:100, BD Biosciences #741319), CD8-eFluor™ 450 (1:200, ThermoFisher #48-0081-82), NK1.1-BV711™ (1:200, BioLegend #108745), CD62L-FITC™ (1:200, ThermoFisher #11-0621082), and CD4-NoaFluor™ Blue 610-70s (1:400, ThermoFisher #M001T02B06-A). Fluorescence minus one samples (FMOs) received all fluorescent antibodies except for one to determine the portion of the spectral signal contributed by that single fluorophore. After 1 hr antibody incubation, the cells were spun down at 1500 RPM for 5 min twice with a 100 μL FACS buffer wash in between. For intracellular staining, cells were fixed in 50 μL fixation/permeabilization solution (eBioscience™ #00-5523-00) overnight at 4°C. Panel intracellular markers included: Ki67-PECy5 (1:200, ThermoFisher #15-5698-82) and FOXP3-Alexa Fluor™ 660 (1:200, ThermoFisher #606-5773-82). The next day, the cells were washed twice with 100 μL permeabilization buffer and centrifuged at 1500 RPM for 5 min. Intracellular staining was performed in 1X Perm buffer for 30-60 min at room temperature, followed by washing with 1X Perm buffer and centrifugation at 1800 RPM for 5 min twice. Cells were resuspended in 200 μL FACS buffer, and 25 μL of CountBright™ Absolute Counting Beads (ThermoFisher #C36950) were added to all samples. Flow cytometry data were acquired using the Cytek Aurora Borealis 5-Laser Flow Cytometer (Cytek Biosciences) and SpectroFlo v.3.0.3 software (Cytek Biosciences). Analysis was performed using FCS Express 7 software (De Novo Software).

### Immunofluorescence

Immunofluorescence methods are described in *Supplemental Methods*.

### Statistical Analysis

Statistical analyses were performed in GraphPad Prism 10 (GraphPad Software). The number of mice per group is reported by “n” value in the figure legends. Results were expressed as means ± SEM. The statistical significance was set at p < 0.05. Two-way analyses of variance (ANOVA) with appropriate multiple comparisons testing were used for comparisons involving more than two groups or for analysis of two groups over time. Welch’s t-test was used for comparisons between two groups when appropriate.

Pearson’s correlation coefficient (r) and the coefficient of determination (R^2^) were calculated to assess relationships between PCD emission data and T-cell percentages or absolute numbers obtained by flow cytometry. Specific statistical tests used for each experiment are detailed in the corresponding figure legends.

## Results

### FUS-mediated BTBO in 3x CRISPR Tumors

Previous work has defined the vascularization kinetics of the 3x CRISPR GEMM glioma model, demonstrating that tumor leakiness emerges between postnatal day 65-80 (P65-80)[2]. Tumor-bearing mice were identified at P30 using in vivo whole-body fluorescent imaging to detect luciferase-expressing tumor cells (**Figures S2**). Tumor presence was confirmed in some mice by H&E staining (**Figure S3**). After P65, tumors were evaluated by magnetic resonance imaging (MRI) to confirm visualization and assess responsiveness to FUS-mediated BTBO. T2-weighted MRI scans, which reflect tissue water content and edema, enabled clear delineation of tumor borders in 3x CRISPR GEMM mice (**Figure 1A**). Baseline contrast-enhanced T1-weighted imaging demonstrated minimal contrast extravasation, consistent with low baseline vascular permeability (**Figure 1B**). Following FUS+MB treatment, T1-weighted imaging revealed a pronounced increase in intratumoral contrast enhancement (**Figure 1C**). Quantification of Multihance™ signal intensity demonstrated a significant fold increase in signal within the tumor relative to the contralateral region of interest after FUS treatment (**Figure 1D**). These results indicate that 3x CRISPR GEMM tumors respond to FUS+MB-mediated BTBO, providing a metric for confirming BTB disruption and assessing changes in vascular permeability in this pathophysiological glioma model.

**Figure 1.**
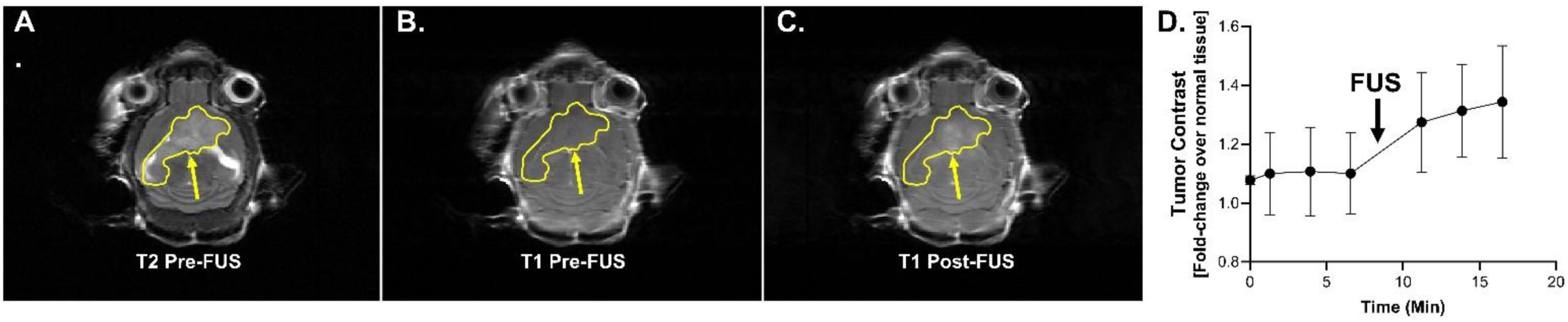
3x CRISPR GEMM are responsive to FUS+MBs. A) Representative T2 MRI image pre-FUS. B) Representative T1 MRI image pre-FUS. C) Representative T1 MRI image post-FUS. D) Fold change of Multihance™ (nM) signal in the tumor over the contralateral side in the 3x CRISPR tumor model over time (minutes). Cyan arrow defines the time of FUS relative to the fold change data.

### 3x CRISPR Tumor-Immune Landscape Remodeling after FUS

Having established that 3x CRISPR GEMM gliomas are responsive to FUS-mediated BBB disruption, we next characterized the tumor-immune landscape three days post-FUS using flow cytometry (**Figure S4**). At baseline, immune cell populations were largely comparable between control and tumor-bearing mice in the absence of FUS+MBs (**Figure 2A-J; Figure S5**). Following FUS, we observed a significant remodeling of the tumor immune compartment. The proportion of total CD45^+^ immune cells increased in the tumor+FUS group compared to tumor alone (p = 0.0411), although absolute CD45^+^ numbers remained unchanged (**Figure 2A, B**). Additionally, the tumor+FUS group exhibited a significantly higher percentage of CD45^+^ cells compared to control+FUS mice (p = 0.0134) (**Figure 2A**).

**Figure 2.**
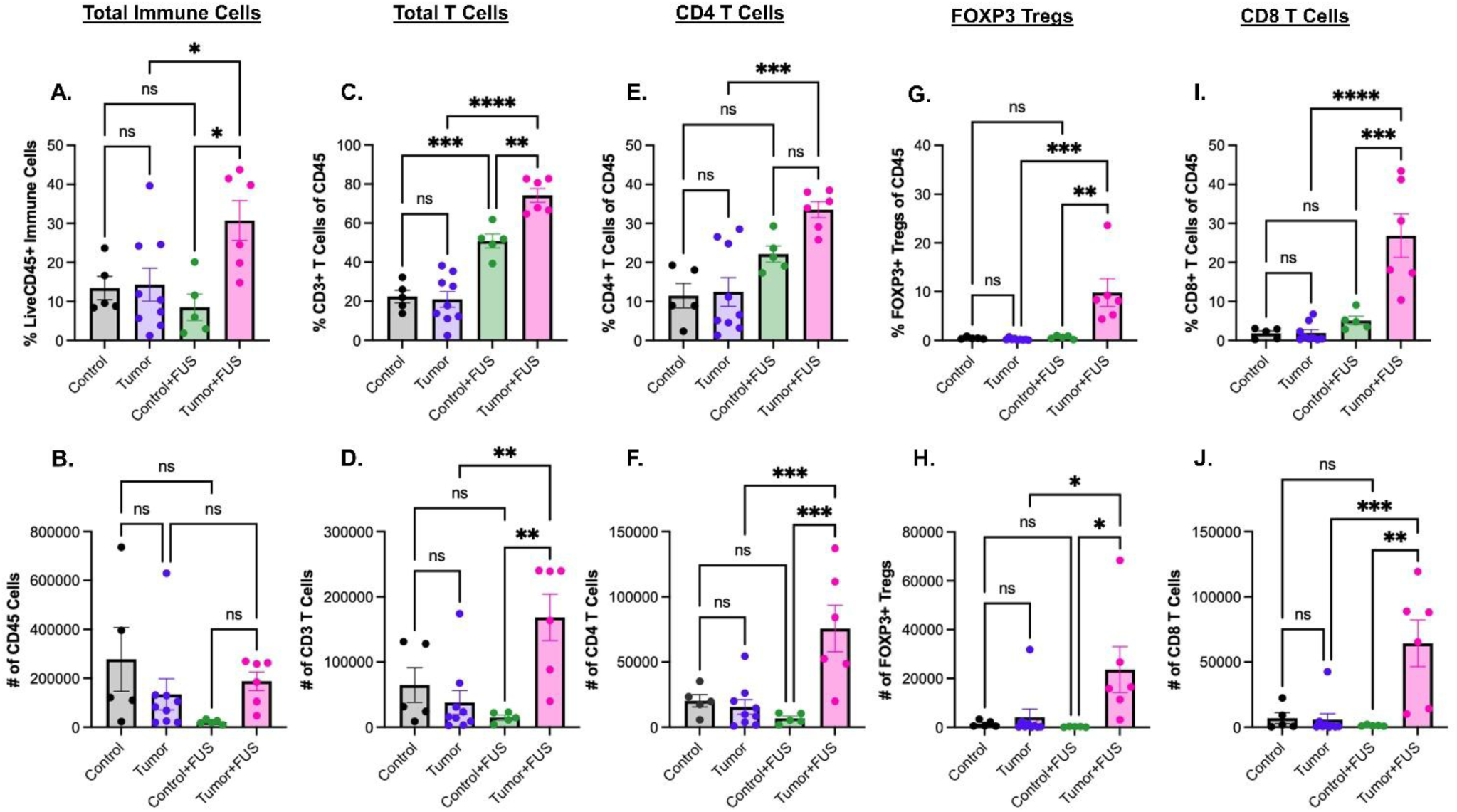
FUS treatment increases brain T-cell populations in 3x CRISPR mice 3 days post-FUS treatment. A) Percent total immune cells (Live/CD45^+^). B) Number of total CD45^+^ immune cells. C) Percent total T cells (Live/CD45^+^CD3^+^). D) Number of total T cells. E) Percent CD4^+^ T cells (Live/CD45^+^CD3^+^CD4^+^). F) Number of CD4^+^ T cells. G) Percent FOXP3^+^ regulatory T cells (Tregs) (Live/CD45^+^CD3^+^CD4^+^FOXP3^+^). H) Number of FOXP3^+^ Tregs. I) Percent CD8^+^ T cells (Live/CD45^+^CD3^+^CD8^+^). J) Number of CD8^+^ T cells. Two-way ANOVA with multiple comparisons correction (Tukey’s). Means ± SEM. * p < 0.05. ** p < 0.01. *** p < 0.001. **** p < 0.0001.

FUS markedly enhanced T-cell infiltration (**Figure S6**). The percentage of CD3^+^ T cells was significantly elevated in FUS-treated groups compared to both control and tumor-only groups, with the tumor+FUS condition showing the greatest increase relative to tumor alone (p < 0.0001), followed by control+FUS (p = 0.0009) (**Figure 2C**). Absolute CD3^+^ T-cell counts rose approximately 4-fold following FUS in tumors (p = 0.0027) and remained 11-fold higher than control+FUS (p = 0.0021) (**Figure 2D**).

Similarly, both the percentage and total number of CD4^+^ T cells were significantly increased in the tumor+FUS group compared to tumor alone and control+FUS groups (**Figure 2E, F**). FOXP3^+^ regulatory T cells (Tregs) were also expanded, with higher counts observed in tumor+FUS mice compared to control+FUS (p = 0.0341) and tumor alone (p = 0.0449) (**Figure 2G, H**). Consistent with this increase, the proportion of FOXP3^+^ Tregs of all CD4^+^ T cells was also elevated suggesting a shift in the tumor inflammatory milieu (**Figure S7**). These findings were supported by representative immunofluorescence images from control, tumor, control+FUS, and tumor+FUS groups, which demonstrated increased CD3/CD4 and CD3/CD8 colocalization in FUS-treated GEMM tumors (**Figure 3**). In addition, T cells appeared more broadly distributed throughout the tumors following FUS treatment, whereas in tumor-only samples, T cells were more frequently clustered near larger vessels (**Figure 3**).

**Figure 3.**
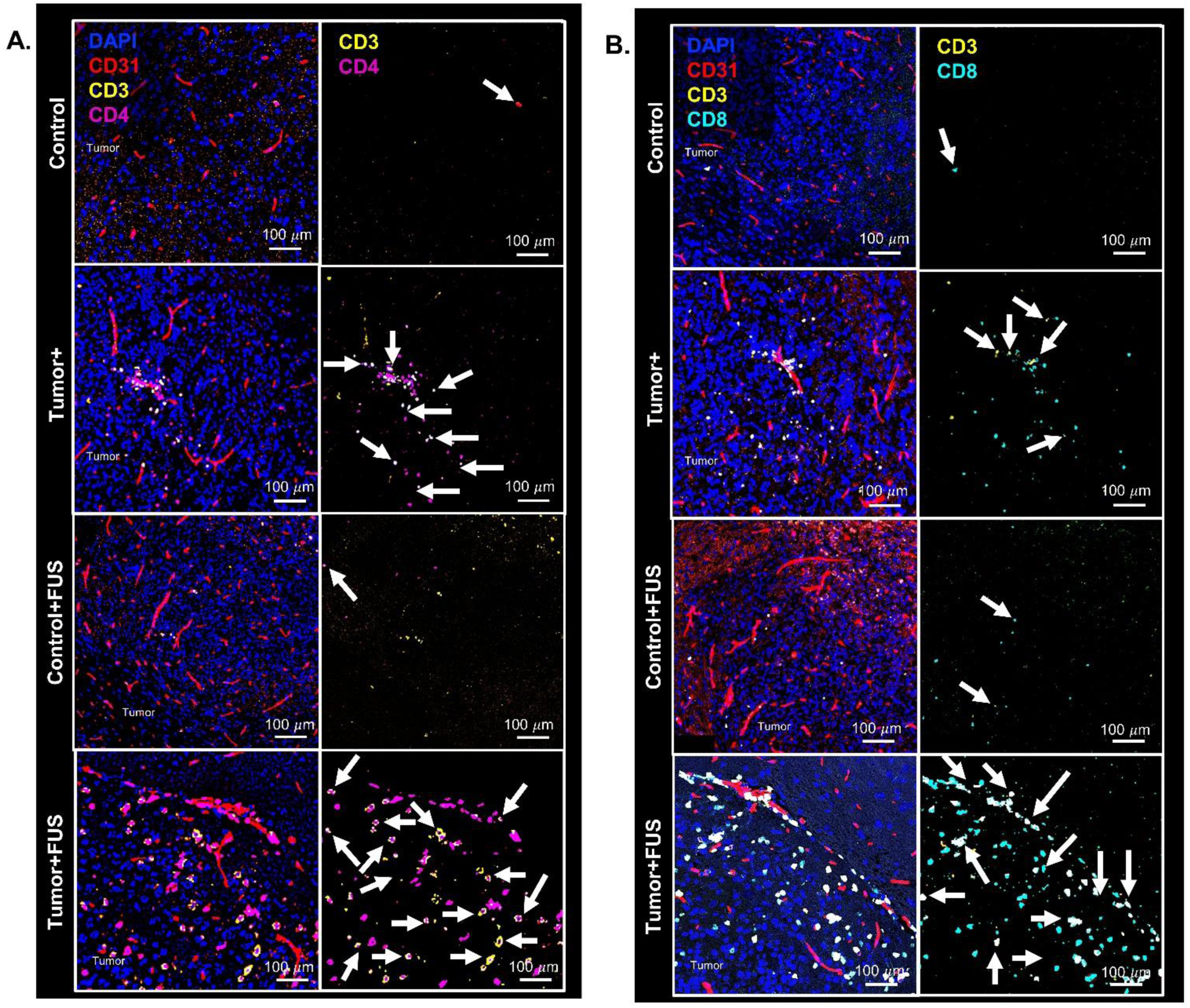
3x CRISPR mice show higher tumor CD4 and CD8 T-cell counts 3 days after FUS. A) Representative immunofluorescent images from control, tumor+, control+FUS, and tumor+FUS mice stained with DAPI (blue), CD31 (red), CD3 (yellow), and CD4 (magenta). Arrows denote colocalization of CD3 with CD4. B) Representative immunofluorescent images from control, tumor+, control+FUS, and tumor+FUS mice stained with DAPI (blue), CD31 (red), CD3 (yellow), and CD8 (cyan). Arrows denote colocalization of CD3 with CD8.

CD8^+^ T cells exhibited the most pronounced response, increasing 5-fold relative to control+FUS (p = 0.0002) and 14-fold relative to tumor alone (p < 0.0001) (**Figure 2I**). Absolute CD8^+^ T-cell numbers were likewise significantly higher in tumor+FUS mice compared to both control+FUS (p = 0.0012) and tumor-only groups (p = 0.0006) (**Figure 2J**). This robust CD8^+^ T-cell accumulation suggests that FUS preferentially enhances cytotoxic T-cell infiltration, consistent with enhanced antitumor immune activation. These findings indicate that FUS broadly amplifies T-cell infiltration while increasing the proportion of CD4^+^ T cells expressing FOXP3^+^ Tregs (**Figure S7**).

In contrast to the pronounced T-cell response, natural killer (NK) cells and myeloid populations exhibited minimal changes across groups (**Figure S5**). Although the fraction of NK cells within the CD45^+^ compartment was higher in tumor+FUS compared to control+FUS (p = 0.0461), absolute NK cell numbers were not significantly different (**Figure S5A, B**). Similarly, the proportions and numbers of dendritic cells (DCs) and CD45^+^CD11b^hi^ monocytes remained stable across all experimental groups (**Figure S5C, D**). Notably, the percentage of CD45^+^CD11b^lo^ microglia was reduced in tumor+FUS compared to both tumor-only (p = 0.0001) and control+FUS groups (p = 0.0089) (**Figure S5E**).

However, when quantified as total counts, microglial numbers were significantly decreased in control+FUS compared to untreated controls (**Figure S5F**). Despite modest shifts in population frequencies in the brain, splenic immune cell subsets remained largely unchanged (**Figure S8**). The exception was the percentage of CD4^+^ T cells, which was significantly increased with FUS in both control and tumor-bearing mice; however, total cell numbers did not change (**Figure S8**). Collectively, these findings demonstrate that FUS+MBs preferentially enhance T-cell infiltration into 3x CRISPR gliomas without broadly altering innate immune populations. The observed accumulation of CD4^+^, CD8^+^, and regulatory T-cell subsets supports the conclusion that FUS-induced BTBO and SI facilitate targeted T-cell recruitment to the TME in this physiologically relevant model.

### Unconventional T-Cell Populations in 3x CRISPR Gliomas

Interestingly, 3x CRISPR gliomas contained distinct populations of unconventional T cells, including CD3^+^CD4^-^CD8^-^ double-negative (DN) and CD3^+^CD4^+^CD8^+^ double-positive (DP) T cells (**Figure 4**). Following FUS+MB treatment, the percentages of both DN and DP T cells increased significantly in control mice (DN, p < 0.0001; DP, p < 0.0001), but not in tumor-bearing mice (**Figure 4A, C**). However, quantification of absolute cell counts revealed no significant differences across groups in either the brain (**Figure 4B, D**) or spleen (**Figure S9**). These findings suggest that 3x CRISPR GEMMs harbor distinct T-cell subsets that may differ from those observed in traditional orthotopic glioma models.

**Figure 4.**
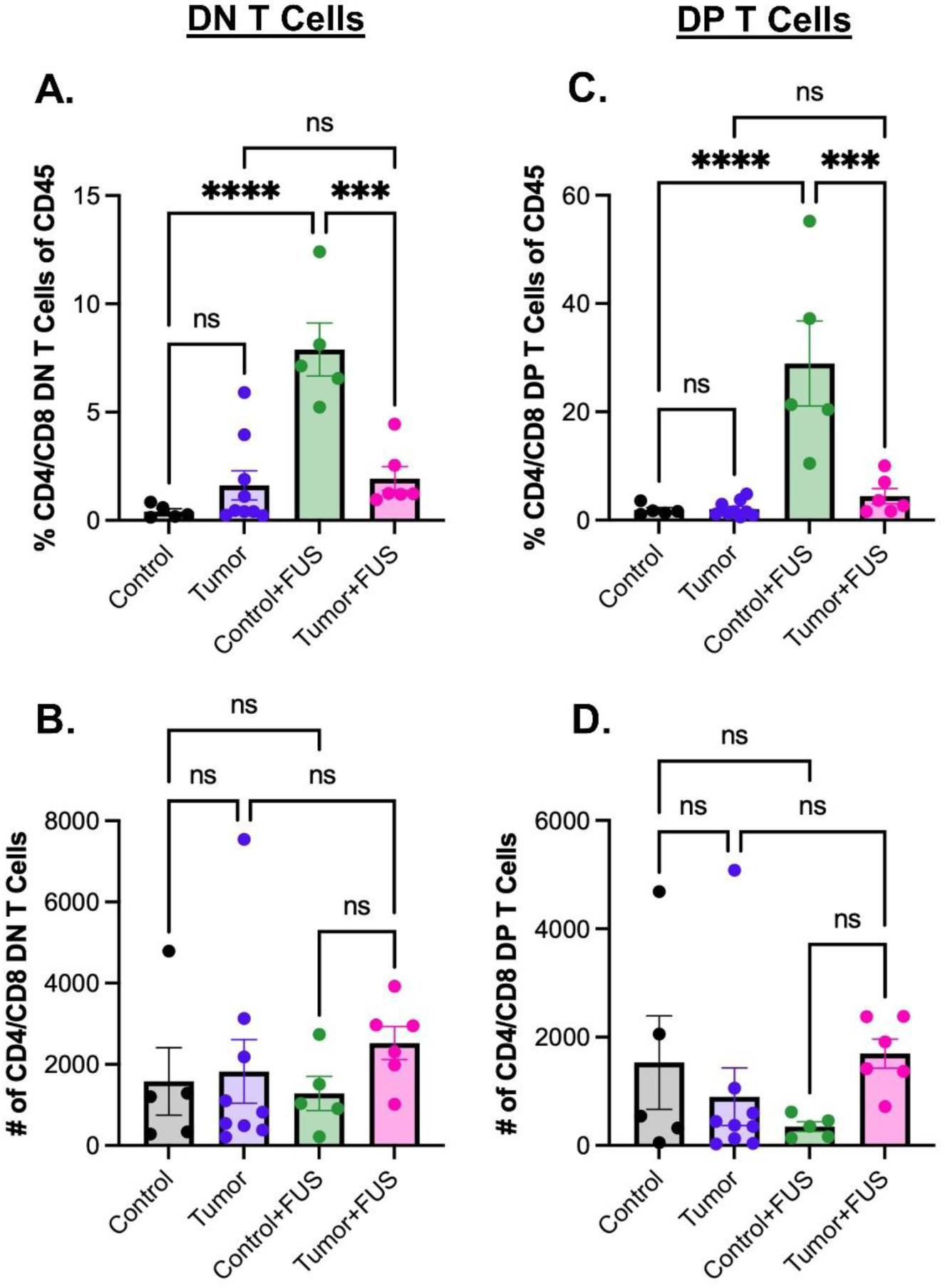
Double negative (DN) and double positive (DP) T cells are present at similar levels in 3x CRISPR GEMM tumors 3 days after FUS. A) Percent double negative (DN; CD4^-^/CD8^-^) T cells in the brain. B) Number of DN T cells. C) Percent double positive (DP; CD4^+^/CD8^+^) T cells in the brain. D) Number of DP T cells. Control (black, n = 5). Tumor (purple, n = 9). Control+FUS (green, n = 5). Tumor+FUS (pink, n = 6). Two-way ANOVA with multiple comparisons correction (Tukey’s). Means ± SEM. * p < 0.05. ** p < 0.01. *** p < 0.001. **** p < 0.0001.

#### Evaluation of Immune Cell Proliferation After FUS+MBs

To determine whether increased immune cell accumulation following FUS+MB treatment was driven by local proliferation or increased infiltration, we assessed the percentage and absolute number of Ki67^+^ immune cells using flow cytometry (**Figure 5**). CD8^+^ T cells, FOXP3^+^ Tregs, and NK cells exhibited no significant differences in Ki67 across experimental groups (**Figure 5C-H**). The percentage of Ki67^+^ CD4 T cells increased 5-fold in the control+FUS group compared to controls (p = 0.0016) (**Figure 5A**); however, this increase was not reflected in total Ki67 CD4^+^ T-cell counts. Instead, tumor+FUS brains exhibited significantly higher numbers of Ki67^+^ CD4^+^ T cells relative to tumor-only mice (p = 0.0373) (**Figure 5B**). In contrast, no significant differences in Ki67 were detected among splenic immune populations (**Figure S10**). Together, these findings suggest that while localized proliferative activity is detectable in select T-cell subsets, the overall increase in T-cell abundance within 3x CRISPR GEMM gliomas following FUS+MBs does not appear to be primarily driven by proliferation.

**Figure 5.**
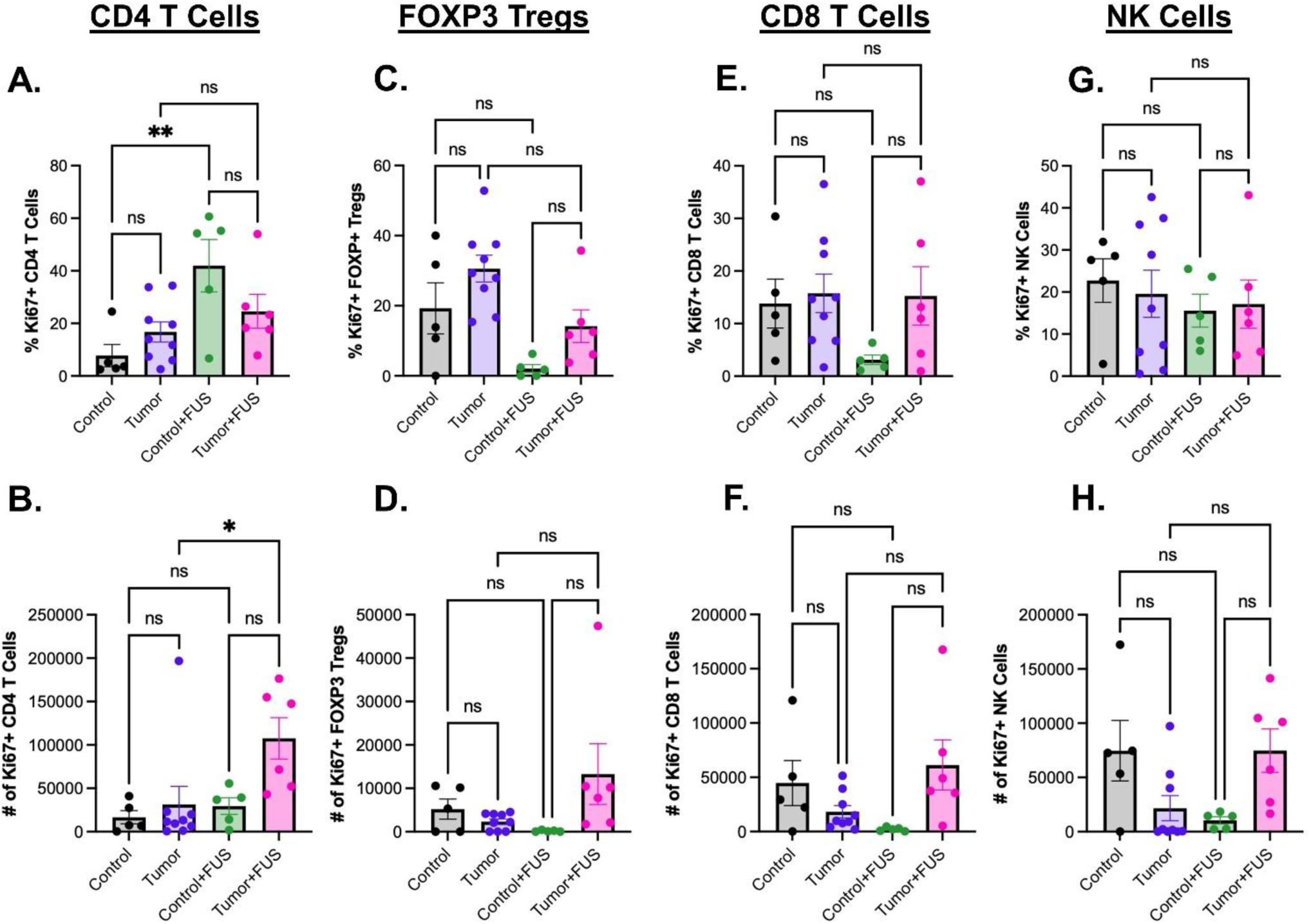
FUS treatment induces minimal changes in the proliferation of brain immune cells in 3x CRISPR mice. A) Percent Ki67^+^ CD4^+^ T cells. B) Number of CD4^+^ T cells. C) Percent Ki67^+^ FOXP3^+^ regulatory T cells (Tregs). D) Number of FOXP3^+^ Tregs. E) Percent Ki67^+^ CD8^+^ T cells. F) Number of CD8^+^ T cells. G) Percent Ki67^+^ natural killer (NK) cells. H) Number of NK cells. Two-way ANOVA with multiple comparisons correction (Tukey’s). Means ± SEM. * p < 0.05. ** p < 0.01.

### Temporal Analysis of FUS-Induced Immune Remodeling

Following our initial analysis at three days post-FUS, we next examined the kinetics of the T-cell response in 3x CRISPR tumors at seven days after FUS treatment (**Figure 6; Figure S11-S16**). The percentage of total live immune cells among all cells was significantly increased in the control+FUS group compared to untreated controls (p = 0.0418); however, absolute CD45^+^ cell numbers were unchanged, suggesting that these differences reflect redistribution among brain and immune cell populations rather than global immune cell expansion (**Figure 6A, B**). No significant differences were observed in major T-cell subsets between tumor and tumor+FUS groups or between the control and control+FUS groups, with one exception: both the percentage and absolute number of FOXP3^+^ Tregs were increased in the control+FUS group relative to control alone (p = 0.0135; p = 0.0507) (**Figure 6 C-J**). All other immune populations remained relatively stable across groups, including in the spleen. We further analyzed effector (CD44^hi^CD62L^lo^), naïve (CD44^lo^CD62L^hi^), and memory (CD44^hi^CD62L^hi^) CD4^+^ and CD8^+^ T-cell subsets within the brain seven days after FUS. Both the percentage and absolute numbers of CD4^+^ and CD8^+^ effector, naïve, and memory T cells were comparable across groups (**Figure S14**). Ki67 expression in T-cell populations was also largely unchanged across groups. The only significant difference was observed in the control versus control+FUS comparison, where FUS increased both the percentage and number of Ki67^+^ CD4^+^ T cells and FOXP3^+^ Tregs (**Figure S15**).

**Figure 6.**
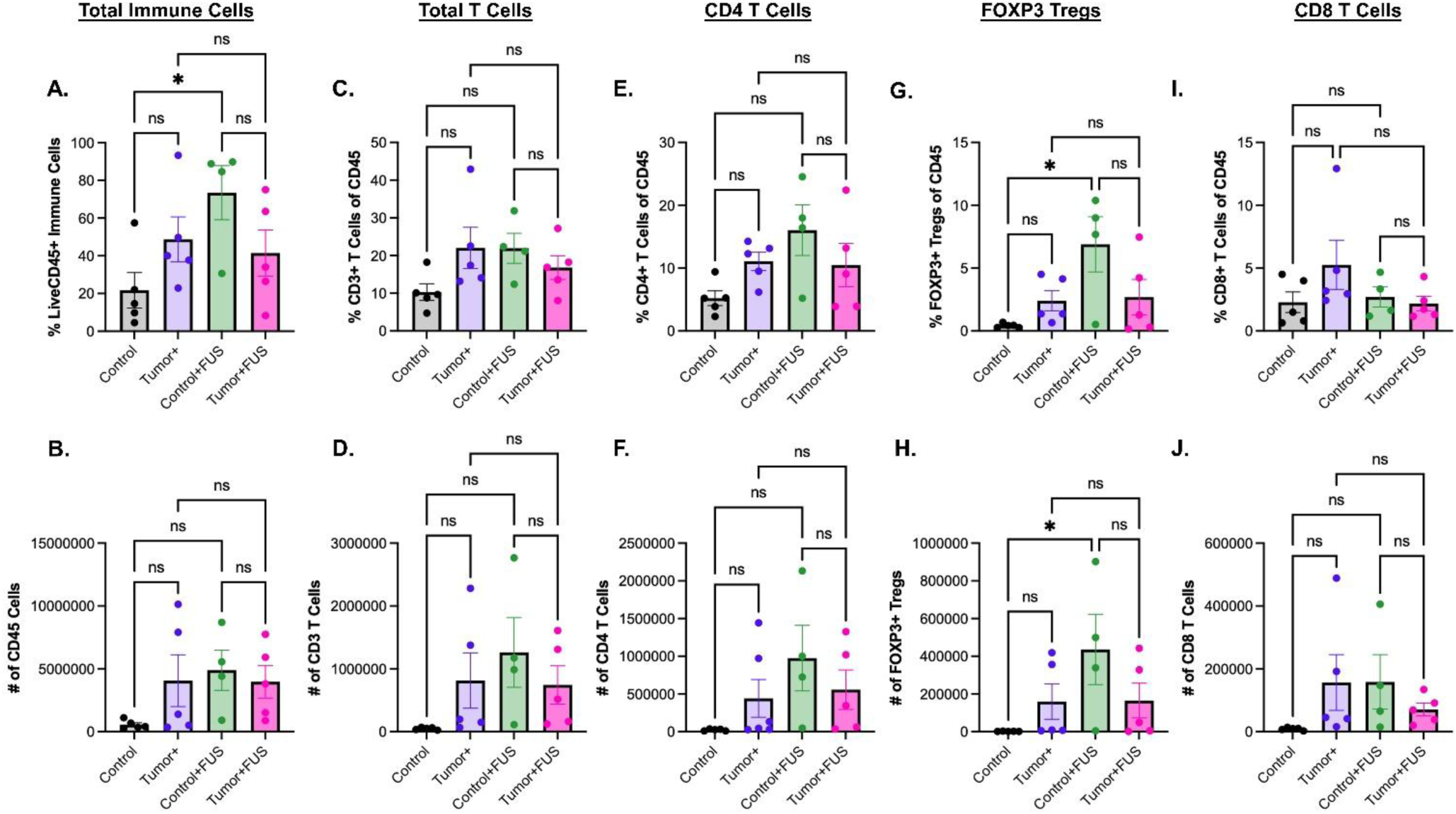
No overall increase in brain T cells was observed 7 days post-FUS in 3x CRISPR mice. A) Percent total immune cells (Live/CD45^+^). B) Number of total CD45^+^ immune cells. C) Percent total T cells (Live/CD45^+^CD3^+^). D) Number of total T cells. E) Percent CD4^+^ T cells (Live/CD45^+^CD3^+^CD4^+^). F) Number of CD4^+^ T cells. G) Percent FOXP3^+^ regulatory T cells (Tregs) (Live/CD45^+^CD3^+^CD4^+^FOXP3^+^). H) Number of FOXP3^+^ Tregs. I) Percent CD8^+^ T cells (Live/CD45^+^CD3^+^CD8^+^). J) Number of CD8^+^ T cells. Two-way ANOVA with multiple comparisons correction (Tukey’s). Means ± SEM. * p < 0.05.

Because the overall immune response to FUS BTB opening at day 7 was remarkably quiescent when compared to day 3, we wanted to confirm that MB oscillation had been successfully driven with FUS. To this end, we examined the PCD data from these experiments, which confirmed that robust acoustic emissions were indeed generated, with no significant differences observed between treatment groups or across time points (**Figure S17**). T-cell populations did not correlate with PCD emissions amplitude (MPa) across FUS-treated groups (**Figure S18**). Collectively, these findings demonstrate that while FUS induces significant immune remodeling at three days post-FUS treatment, the associated T-cell expansion is not sustained at seven days. This suggests that FUS-mediated immune activation is transient and may define a limited, optimal therapeutic window for modulating the local tumor immune environment.

## Discussion

In addition to facilitating drug delivery to GBM, FUS BTB opening has the potential to generate a sterile inflammatory response that opposes the baseline immunosuppressive microenvironment. However, studies examining this response in mice have been limited to implanted tumor models that do not fully recapitulate the BTB, infiltrative growth patterns, or immune landscape of human GBM[26].

Herein, we demonstrate that FUS+MB-mediated BTB disruption yields an unexpectedly robust augmentation of adaptive immune infiltration in a physiologically relevant 3x CRISPR GEMM of glioma. While tumor presence alone induced minimal immune remodeling, FUS treatment significantly increased intratumoral CD4^+^, CD8^+^, and FOXP3^+^ regulatory T-cell representation. Importantly, increases in these cell populations were not accompanied by marked elevations in proliferation, consistent with the hypothesis that FUS BTB opening enhances immune cell recruitment, rather than stimulating local cell proliferation. The ratio of FOXP3^+^ Tregs to CD4^+^ T cells was increased in tumors following FUS, suggesting that BTBO amplifies T-cell recruitment while also altering the inflammatory immune milieu. In contrast, innate immune populations were largely unchanged, supporting a model in which FUS preferentially modulates adaptive immune accessibility, possibly through increased integrin or chemokine expression by endothelial cells. Collectively, the results suggest that FUS BTB opening could exert an unexpectedly pronounced impact on the recruitment of adoptive T-cell therapies to GBM.

Previous studies have reported acute inflammatory responses following FUS BBB disruption, including enhanced antigen presentation, endothelial activation, microglial proliferation, and monocyte or DC recruitment in both naïve and tumor-bearing brains[27–29]. Additionally, FUS+MB treatment facilitates trafficking of adoptively transferred CAR T cells to intracranial tumors[11]. However, our study provides the first evidence of altered endogenous adaptive immune populations following FUS-mediated BTB disruption using clinically relevant FUS parameters (**Figure 2, 3**). Previous studies in implantable tumor models have generally reported no change in T-cell frequency or number in GBM tumors, or have required high mechanical indices to observe increased T-cell infiltration[29–31]. In contrast, our data in a native, immunocompetent mouse model of glioma demonstrate unexpectedly robust recruitment of T cells to the tumor microenvironment after FUS. The most pronounced effects were observed in CD8^+^ T cells, which increased up to 14-fold in tumor+FUS mice compared to tumor-only controls (**Figure 2**). This bias towards cytotoxic T-cell infiltration is notable, as CD8^+^ T cells are central mediators of anti-tumor immunity. These data suggest that FUS-induced vascular modulation may selectively facilitate the trafficking of tumor-reactive cytotoxic T cells, raising the possibility that FUS could prime gliomas for combination immunotherapies such as checkpoint blockade or adoptive T-cell transfer.

T cells in tumor-only samples appeared to cluster around larger blood vessels, whereas FUS treatment was associated with a broader distribution of T cells throughout the tumor (**Figure 3**). This observation is consistent with previous reports that T cells frequently localize near the vasculature in gliomas, particularly in IDH-mutant tumors[32,33]. In other cancer types, the spatial organization of T cells has been linked to therapeutic outcomes. For example, in colorectal cancer, increased perivascular T-cell localization has been associated with improved responses to immunotherapy, whereas in melanoma, a more diffuse intratumoral distribution of T cells has been correlated with improved survival[34,35]. Together, these findings suggest that FUS-mediated vascular modulation may influence not only the magnitude but also the spatial distribution of tumor-infiltrating T cells. Such changes in T-cell localization could potentially enhance antitumor immune activity and improve responsiveness to immunotherapies in glioma.

Although we observed a modest increase in the percentage of CD4^+^ T cells in the spleen three days after FUS in both control and tumor-bearing mice, this splenic change was only ∼1.5-fold, compared with a ∼3-fold increase in the brain. Thus, the systemic splenic response is insufficient to account for the pronounced T-cell accumulation in the TME. This observation may reflect local effects of MB accumulation in the spleen during circulation, which could transiently influence CD4+ T-cell distribution without driving the robust infiltration observed in the brain. Collectively, these results indicate that FUS-induced modulation of adaptive immunity is primarily localized to the brain, highlighting the potential of FUS to reshape the TME without inducing widespread systemic effects.

Temporal analysis revealed that the enhanced T-cell infiltration observed at three days post-FUS was largely transient, with minimal differences between tumor and tumor+FUS groups by seven days (**Figure 6**). This transient response likely reflects the temporary nature of FUS-mediated BTB opening, which may transiently increase vascular permeability, endothelial activation, and chemokine expression, creating a permissive window for immune cell entry. These findings highlight a critical opportunity to synchronize FUS with immunotherapy administration, suggesting that timed delivery of adoptive T-cell therapies or other immune modulators during this window could maximize tumor infiltration and therapeutic efficacy. Moreover, the stage-dependent vascular characteristics of the 3x CRISPR model likely contribute to these dynamics, as the degree of BTB permeability and vascular remodeling directly influence immune cell trafficking. These findings underscore the importance of vascular context in shaping immune accessibility and may help explain why implantable tumor models, which often exhibit higher baseline BTB permeability, have reported minimal T-cell recruitment following FUS[29–31].

We identified distinct populations of unconventional T-cell populations, including CD3^+^CD4^-^CD8^-^DN and CD3^+^CD4^+^CD8^+^ DP T cells, within the 3x CRISPR gliomas (**Figure 4**). DN T cells, often γδ T cell receptor-expressing or NK-T cells, have been described in multiple solid tumors and can exert both pro-and anti-tumor functions[36–41]. These cells have been detected in patient-derived glioma tissue as well as in preclinical GBM models, suggesting a conserved role in the glioma immune landscape[37–39]. In contrast, DP T cells—rare in both humans and mice—are typically found in peripheral blood and lymphoid tissues and have been reported in select pathological conditions, including cancer[40,42]. To our knowledge, this is the first report of DP T cells in a glioma model.

Ki67 analysis indicated minimal changes in T-cell proliferation following FUS, suggesting that the observed increases in T-cell abundance reflects enhanced trafficking rather than local expansion (**Figure 5**). This aligns with the suppressive metabolic environment of the glioma TME, characterized by hypoxia, nutrient depletion, and accumulation of immunosuppressive metabolites, which may limit local proliferation of effector T cells while favoring Treg persistence[43,44]. Future metabolomic profiling of these microenvironments will be important for understanding how FUS influences nutrient availability, T-cell metabolism, and functional capacity within the TME.

Collectively, these results establish that FUS+MB-mediated BTBO transiently remodels the adaptive immune landscape in a physiologically relevant glioma model without broadly altering innate compartments. The selective recruitment of CD8 T cells and maintenance of immune homeostasis suggest that FUS can convert an otherwise immunologically restrictive microenvironment into a window of enhanced immune accessibility. These findings provide a foundation for leveraging FUS-mediated SI to improve immunotherapeutic efficacy in GBM.

## Acknowledgments

Supported by NIH R01CA279134, R21CA286367, and R01NS111102 to RJP. KMN was supported by a training fellowship from the UVA Comprehensive Cancer Center and an NIH Training Grant (NS115657) to THH. MRI was performed in the University of Virginia Molecular Imaging Core Laboratory, with support for the 9.4T Bruker scanner from NIH S10OD025024. Flow Cytometry was performed in the University of Virginia Flow Cytometry Core. Immunofluorescence was performed by the Biorepository and Tissue Research Facility, which is supported by the University of Virginia School of Medicine, Research Resource Identifiers (RRID): SCR_022971.

## Data and Materials Availability

All data are provided in the manuscript. All plasmids used to generate the 3x CRISPR tumor model have been deposited at Addgene (https://www.addgene.org/Joshua_Wythe/).

## Author Contributions

Conceptualization: KMN, MRH, THH, JDW, RJP Formal

Analysis: KMN, MRH, TRB

Funding Acquisition: KMN, THH, JDW, RJP

Methodology: KMN, MRH, CAC, WBG, VRB, ENG, TC, CMG

Investigation: KMN, MRH, CAC, WBG, VRB, ENG, TC, CMG

Supervision: THH, JDW, RJP

Writing—original draft: KMN Writing—review & editing: KMN, MRH, CAC, WBG, VRB, ENG, TC, CMG, THH, JDW, RJP

## References

[1] R.K. Jain, E. di Tomaso, D.G. Duda, J.S. Loeffler, A.G. Sorensen, T.T. Batchelor, Angiogenesis in brain tumours, Nat. Rev. Neurosci. 8 (2007) 610–622. 10.1038/nrn2175.

2. J.C. Carlson, M. Cantu Gutierrez, B. Lozzi, E. Huang-Hobbs, W.D. Turner, B. Tepe, Y. Zhang, A.M. Herman, G. Rao, C.J. Creighton, J.D. Wythe, B. Deneen, Identification of diverse tumor endothelial cell populations in malignant glioma, Neuro. Oncol. 23 (2020) 932–944. 10.1093/neuonc/noaa297.

[3] K. Singh, K.M. Hotchkiss, K.K. Patel, D.S. Wilkinson, A.A. Mohan, S.L. Cook, J.H. Sampson, Enhancing T Cell Chemotaxis and Infiltration in Glioblastoma, Cancers (Basel). 13 (2021) 5367. 10.3390/cancers13215367.

[4] Z.I. Kovacs, S. Kim, N. Jikaria, F. Qureshi, B. Milo, B.K. Lewis, M. Bresler, S.R. Burks, J.A. Frank, Disrupting the blood-brain barrier by focused ultrasound induces sterile inflammation., Proc. Natl. Acad. Sci. U. S. A. 114 (2017) E75–E84. 10.1073/pnas.1614777114.

[5] C. Kim, M. Lim, G.F. Woodworth, C.D. Arvanitis, The roles of thermal and mechanical stress in focused ultrasound-mediated immunomodulation and immunotherapy for central nervous system tumors, J. Neurooncol. (2022). 10.1007/s11060-022-03973-1.

[6] P.J. Martinez, J.J. Song, J.I. Castillo, J. DeSisto, K.-H. Song, A.L. Green, M. Borden, Effect of Microbubble Size, Composition, and Multiple Sonication Points on Sterile Inflammatory Response in Focused Ultrasound-Mediated Blood–Brain Barrier Opening, ACS Biomater. Sci. Eng. 10 (2024) 7451–7465. 10.1021/acsbiomaterials.4c00777.

[7] A.R. Kline-Schoder, S. Chintamen, M.J. Willner, M.R. DiBenedetto, R.L. Noel, A.J. Batts, N. Kwon, S. Zacharoulis, C.C. Wu, V. Menon, S.G. Kernie, E.E. Konofagou, Characterization of the responses of brain macrophages to focused ultrasound-mediated blood–brain barrier opening, Nat. Biomed. Eng. 8 (2024) 650–663. 10.1038/s41551-023-01107-0.

[8] K.L. Rock, E. Latz, F. Ontiveros, H. Kono, The sterile inflammatory response, Annu. Rev. Immunol. 28 (2010) 321–342. 10.1146/annurev-immunol-030409-101311.

[9] A.S. Mathew, C.M. Gorick, E.A. Thim, W.J. Garrison, A.L. Klibanov, G.W. Miller, N.D. Sheybani, R.J. Price, Transcriptomic response of brain tissue to focused ultrasound-mediated blood–brain barrier disruption depends strongly on anesthesia, Bioeng. Transl. Med. 6 (2020) e10198. 10.1002/btm2.10198.

[10] C. Angolano, E. Hansen, H. Ajjawi, P. Nowlin, Y. Zhang, N. Thunemann, C. Ferran, N. Todd, Characterization of focused ultrasound blood-brain barrier disruption effect on inflammation as a function of treatment parameters, Biomedicine & Pharmacotherapy 182 (2025) 117762. 10.1016/j.biopha.2024.117762.

[11] Y. Guo, H. Lee, C. Kim, C. Park, A. Yamamichi, P. Chuntova, M. Gallus, M.O. Bernabeu, H. Okada, H. Jo, C. Arvanitis, Ultrasound frequency-controlled microbubble dynamics in brain vessels regulate the enrichment of inflammatory pathways in the blood-brain barrier, Nat. Commun. 15 (2024) 8021. 10.1038/s41467-024-52329-y.

[12] D. McMahon, A. Lassus, E. Gaud, V. Jeannot, K. Hynynen, Microbubble formulation influences inflammatory response to focused ultrasound exposure in the brain, Sci. Rep. 10 (2020) 21534. 10.1038/s41598-020-78657-9.

[13] D. McMahon, K. Hynynen, Acute Inflammatory Response Following Increased Blood-Brain Barrier Permeability Induced by Focused Ultrasound is Dependent on Microbubble Dose, Theranostics 7 (2017) 3989–4000. 10.7150/thno.21630.

14. S.D. Finkelstein, P. Black, T.P. Nowak, C.M. Hand, S. Christensen, P.W. Finch, Histological Characteristics and Expression of Acidic and Basic Fibroblast Growth Factor Genes in Intracerebral Xenogeneic Transplants of Human Glioma Cells, Neurosurgery 34 (1994) 136. https://journals.lww.com/neurosurgery/fulltext/1994/01000/histological_characteristics_and_expression_of.20.aspx.

[15] W. Maes, S.W. Van Gool, Experimental immunotherapy for malignant glioma: lessons from two decades of research in the GL261 model, Cancer Immunology, Immunotherapy 60 (2011) 153–160. 10.1007/s00262-010-0946-6.

[16] K.B. Grausam, J.J. Breunig, Modeling Brain Tumors In Vivo Using Electroporation-Based Delivery of Plasmid DNA Representing Patient Mutation Signatures., Journal of Visualized Experiments (2023). 10.3791/65286.

[17] J.K. Khalsa, N. Cheng, J. Keegan, A. Chaudry, J. Driver, W.L. Bi, J. Lederer, K. Shah, Immune phenotyping of diverse syngeneic murine brain tumors identifies immunologically distinct types, Nat. Commun. 11 (2020) 3912. 10.1038/s41467-020-17704-5.

[18] K. Lenting, R. Verhaak, M. ter Laan, P. Wesseling, W. Leenders, Glioma: experimental models and reality, Acta Neuropathol. 133 (2017) 263–282. 10.1007/s00401-017-1671-4.

[19] D. Hambardzumyan, L.F. Parada, E.C. Holland, A. Charest, Genetic modeling of gliomas in mice: new tools to tackle old problems, Glia 59 (2011) 1155–1168. 10.1002/GLIA.21142.

[20] L. Eklund, M. Bry, K. Alitalo, Mouse models for studying angiogenesis and lymphangiogenesis in cancer, Mol. Oncol. 7 (2013) 259–282. 10.1016/j.molonc.2013.02.007.

[21] P.C. Huszthy, I. Daphu, S.P. Niclou, D. Stieber, J.M. Nigro, P.Ø. Sakariassen, H. Miletic, F. Thorsen, R. Bjerkvig, In vivo models of primary brain tumors: pitfalls and perspectives, Neuro. Oncol. 14 (2012) 979–993. 10.1093/neuonc/nos135.

[22] J.A. Nagy, S.-H. Chang, S.-C. Shih, A.M. Dvorak, H.F. Dvorak, Heterogeneity of the Tumor Vasculature, Semin. Thromb. Hemost. 36 (2010) 321–331. 10.1055/s-0030-1253454.

[23] Y. Zhang, S. Wang, A.C. Dudley, Models and molecular mechanisms of blood vessel co-option by cancer cells, Angiogenesis 23 (2020) 17–25. 10.1007/s10456-019-09684-y.

[24] L.A. Genovesi, S. Puttick, A. Millar, M. Kojic, P. Ji, A.K. Lagendijk, C. Brighi, C.S. Bonder, C. Adolphe, B.J. Wainwright, Patient-derived orthotopic xenograft models of medulloblastoma lack a functional blood-brain barrier, Neuro. Oncol. 23 (2021) 732–742. 10.1093/neuonc/noaa266.

[25] W.B. Gillespie III, Y. Zhang, O.E. Ruiz, J. Cerda III, J. Ortiz-Guzman, M. Sherman, W.D. Turner, G. Largoza, L.E. Mosser, E. Fujimoto, C.-B. Chien, K.M. Kwan, B.R. Arenkiel, W.P. Devine, J.D. Wythe, MultiSite Assembly of Gateway Induced Clones (MAGIC): a flexible cloning toolbox for use in vertebrate model systems, Development 152 (2025) dev204308. 10.1242/dev.204308.

[26] N. Yadav, B.W. Purow, Understanding current experimental models of glioblastoma-brain microenvironment interactions, J. Neurooncol. 166 (2024) 213–229. 10.1007/s11060-023-04536-8.

[27] C.T. Curley, A.D. Stevens, A.S. Mathew, K. Stasiak, W.J. Garrison, G. Wilson Miller, N.D. Sheybani, V.H. Engelhard, T.N.J. Bullock, R.J. Price, Immunomodulation of intracranial melanoma in response to blood-tumor barrier opening with focused ultrasound, Theranostics 10 (2020) 8821–8833. 10.7150/thno.47983.

[28] Y. Zhang, J. Wang, S.N. Ghobadi, H. Zhou, A. Huang, M. Gerosa, Q. Hou, O. Keunen, A. Golebiewska, F.G. Habte, G.A. Grant, R. Paulmurugan, K.S. Lee, M. Wintermark, Molecular Identity Changes of Tumor-Associated Macrophages and Microglia After Magnetic Resonance Imaging–Guided Focused Ultrasound–Induced Blood–Brain Barrier Opening in a Mouse Glioblastoma Model, Ultrasound Med. Biol. (2023). 10.1016/J.ULTRASMEDBIO.2022.12.006.

[29] N.D. Sheybani, A.R. Witter, W.J. Garrison, G.W. Miller, R.J. Price, T.N.J. Bullock, Profiling of the immune landscape in murine glioblastoma following blood brain/tumor barrier disruption with MR image-guided focused ultrasound, J Neurooncol 156 (2022) 109–122. 10.1007/s11060-021-03887-4.

[30] K.T. Chen, W.Y. Chai, Y.J. Lin, C.J. Lin, P.Y. Chen, H.C. Tsai, C.Y. Huang, J.S. Kuo, H.L. Liu, K.C. Wei, Neuronavigation-guided focused ultrasound for transcranial blood-brain barrier opening and immunostimulation in brain tumors, Sci. Adv. 7 (2021). 10.1126/SCIADV.ABD0772/SUPPL_FILE/ABD0772_SM.PDF.

[31] K.M. Nowak, M.R. Hoch, V.R. Breza, C.M. Gorick, J. Song, A.C. Debski, J.D. Samuels, M.R. DeWitt, B.W. Purow, T.N. Bullock, T.H. Harris, R.J. Price, The Influence of VEGFR-2 Blockade and Focused Ultrasound (FUS) Blood-Brain Barrier Opening on the Glioma-Immune Landscape, Neurooncol. Adv. (2025) vdaf221. 10.1093/NOAJNL/VDAF221.

[32] M. Dinevska, S.S. Widodo, L. Furst, L. Cuzcano, Y. Fang, S. Mangiola, P.J. Neeson, P.K. Darcy, R.G. Ramsay, R. Hutchinson, F. MacKay, M. Christie, S.S. Stylli, T. Mantamadiotis, Cell signaling activation and extracellular matrix remodeling underpin glioma tumor microenvironment heterogeneity and organization, Cellular Oncology 46 (2023) 589–602. 10.1007/s13402-022-00763-9.

[33] R. Luning, P.J. French, W.J.F. Vanbilloen, L. Dobber, L. Van Hijfte, R. Hoogendijk, M.J. van den Bent, R. Debets, M. Geurts, T cell immunity in glioma and potential implications for immunotherapy: A systematic review, Neuro. Oncol. (2025) noaf236. 10.1093/neuonc/noaf236.

[34] C.R. Stoltzfus, R. Sivakumar, L. Kunz, B.E. Olin Pope, E. Menietti, D. Speziale, R. Adelfio, M. Bacac, S. Colombetti, M. Perro, M.Y. Gerner, Multi-Parameter Quantitative Imaging of Tumor Microenvironments Reveals Perivascular Immune Niches Associated With Anti-Tumor Immunity, Front. Immunol. 12 (2021). 10.3389/fimmu.2021.726492.

[35] G. Erdag, J.T. Schaefer, M.E. Smolkin, D.H. Deacon, S.M. Shea, L.T. Dengel, J.W. Patterson, C.L. Slingluff Jr, Immunotype and Immunohistologic Characteristics of Tumor-Infiltrating Immune Cells Are Associated with Clinical Outcome in Metastatic Melanoma, Cancer Res. 72 (2012) 1070–1080. 10.1158/0008-5472.CAN-11-3218.

[36] T. Ahmedna, H. Khela, C. Weber-Levine, T.D. Azad, C.M. Jackson, K. Gabrielson, C. Bettegowda, J. Rincon-Torroella, The Role of γδ T-Lymphocytes in Glioblastoma: Current Trends and Future Directions, Cancers (Basel). 15 (2023) 5784. 10.3390/cancers15245784.

[37] Z. Wu, Y. Zheng, J. Sheng, Y. Han, Y. Yang, H. Pan, J. Yao, CD3+CD4-CD8- (Double-Negative) T Cells in Inflammation, Immune Disorders and Cancer, Front. Immunol. 13 (2022). 10.3389/fimmu.2022.816005.

[38] F. Galvez-Cancino, M. Navarrete, G. Beattie, S. Puccio, E. Conde-Gallastegi, K. Foster, Y. Morris, T. Sahwangarrom, D. Karagianni, J. Liu, A.J.X. Lee, D.A. Garyfallos, A.P. Simpson, G.-T. Mastrokalos, F. Nannini, C. Costoya, V. Anantharam, B.C. Cianciotti, L. Bradley, C. Garcia-Diaz, M. Clements, A. Shroff, F.V. Dastjerdi, E.M. Rota, S. Sheraz, R. Bentham, I. Uddin, H. Walczak, A. Lladser, J.L. Reading, K.A. Chester, M.A. Pule, P.M. Brennan, S. Marguerat, S. Parrinello, K.S. Peggs, N. McGranahan, E. Lugli, K. Litchfield, S.M. Pollard, S.A. Quezada, Regulatory T cell depletion promotes myeloid cell activation and glioblastoma response to anti-PD1 and tumor-targeting antibodies, Immunity 58 (2025) 1236–1253.e8. 10.1016/j.immuni.2025.03.021.

[39] Z. Liu, Q. Meng, J. Bartek Jr, T. Poiret, O. Persson, L. Rane, E. Rangelova, C. Illies, I.H. Peredo, X. Luo, M.V. Rao, R.A. Robertson, E. Dodoo, M. Maeurer, Tumor-infiltrating lymphocytes (TILs) from patients with glioma, Oncoimmunology 6 (2017) e1252894. 10.1080/2162402X.2016.1252894.

[40] N.H. Overgaard, J.-W. Jung, R.J. Steptoe, J.W. Wells, CD4+/CD8+ double-positive T cells: more than just a developmental stage?, J. Leukoc. Biol. 97 (2015) 31–38. 10.1189/jlb.1RU0814-382.

[41] M. McDonald, Y. Yang, A. Puentes, H. Najem, K. Latha, E. Goethe, Y. Ko, A. Heimberger, A. Harmanci, G. Rao, IMMU-35. CD3+CD4−CD8− (Double-Negative) T-cells Provide Inflammatory Signals That Influence Myeloid Cells and Enhance Survival in Glioblastoma, Neuro. Oncol. 27 (2025) v210. 10.1093/neuonc/noaf201.0833.

[42] J. Desfrançois, A. Moreau-Aubry, V. Vignard, Y. Godet, A. Khammari, B. Dréno, F. Jotereau, N. Gervois, Double Positive CD4CD8 αβ T Cells: A New Tumor-Reactive Population in Human Melanomas, PLoS One 5 (2010) e8437. 10.1371/journal.pone.0008437.

[43] R.D. Leone, J.D. Powell, Metabolism of immune cells in cancer, Nat. Rev. Cancer 20 (2020) 516–531. 10.1038/s41568-020-0273-y.

[44] M. Jiang, H. Fang, H. Tian, Metabolism of cancer cells and immune cells in the initiation, progression, and metastasis of cancer, Theranostics 15 (2025) 155–188. 10.7150/thno.103376.

