## Supplemental Materials for "Focused Ultrasound Blood-Tumor Barrier Opening Rapidly Augments Intratumor CD4 and CD8 T Cell Representation in a Genetically Engineered Mouse Model of Glioma"

### Focused Ultrasound Blood-Brain Barrier Opening Drives a Rapid Increase in Intratumor CD4 and CD8 T Cells in a Genetically Engineered Mouse Model of Glioma

Katherine M. Nowak[1], Matthew R. Hoch[2], William B. Gillespie III[3], Claire A. Conarroe[4], Victoria R. Breza[2], Tanya Cruz[2], Eden N. Gordon[2], Catherine M. Gorick[2], Tajie H. Harris[5], Joshua D. Wythe[3,5], and Richard J. Price[2,6].

1. Department of Microbiology, Immunology, and Cancer Biology, University of Virginia, Charlottesville, VA
2. Department of Biomedical Engineering, University of Virginia, Charlottesville, VA
3. Department of Cell and Developmental Biology, University of Virginia, Charlottesville, VA
4. Department of Pathology, University of Virginia, Charlottesville, VA
5. Department of Neuroscience, University of Virginia, Charlottesville, VA
6. Radiology and Medical Imaging, University of Virginia, Charlottesville, VA

Corresponding Author:
Richard J. Price, Ph.D.
415 Lane Rd, MR5 Building, Room 2316
Charlottesville, VA 22903


#### Supplemental Methods

##### Genetically Engineered Mouse Model of Glioma

Noon on the day a vaginal plug was discovered was defined as E0.5; birth was P0. Adult mice were at least 8 weeks old and housed with access to food (normal chow diet) and water ad libitum on a 12-h light–12-h dark cycle at 21 °C and 50–60% humidity. Timed matings were set up at the end of the workday. For in utero electroporation, pregnant dams were anesthetized and the uterine horns exposed. Six 33 V pulses (50 ms pulse, 75 ms interval) were delivered using a NEPA21 Electro-Kinetic Transfection System (Bulldog Bio). Uterine horns were then placed back in the dam, and the peritoneal muscle was closed using absorbable sutures (McKesson, 99420), and the skin incision was closed with non-absorbable sutures (McKesson, 2685). Dams recovered on a warming disk.

##### MRI Acquisition & Data Processing

MR images (pre- and post-contrast, including post-FUS) were acquired using two sets of spin-echo images (Bruker: RARE) (repetition time = 1040 ms, echo time = 6.71 ms, slice thickness = 0.6 mm, slice gap = 0.6 mm, field of view = 35 × 35 mm, matrix size = 200 × 200, spatial resolution = 0.175 × 0.175 × 0.6 mm$^3$, rare factor = 10, and number of slices = 7). The scans were acquired in an interchanging manner, with the second scan positioned in the slice gap of the first scan to allow for 3D visualization of the tumor. This approach was utilized to minimize crosstalk between adjacent slices. The pre-contrast scan was acquired with one average, and subsequent scans were acquired with four averages, resulting in total scan times of 19.76 seconds and 79.04 seconds, respectively. A total of 11 post-contrast scans were acquired, 6 scans in slice package 1 and 5 scans in slice package 2. FUS was applied after the fifth post-contrast acquisition, with imaging resuming immediately after FUS. MultiHance™ was intravenously injected as a bolus dose of 357 µmol/kg after the pre-contrast agent scans.

A systematic signal decay over time was observed across identical scans. To account for this, we used a custom MATLAB script to manually define ROIs around the tumor in slices where it was visible and within a contralateral, tumor-free brain region. The average receiver gain–normalized signal within the tumor and contralateral ROIs were then ratioed to normalize to each scan's specific decay and calculate fold changes.

**Passive Cavitation Detection & Data Processing**

PCD emissions were collected during FUS treatment using the FUS Aureus software (FUS Instruments). Baseline emissions were obtained using 20 sonications at 0.325 MPa without MBs. PCD files were processed with a custom MATLAB script (MathWorks) to extract the PNP and quantify PCD emissions over the FUS+MB treatment period for each sonication target. Emissions were quantified as AUC with a 300 Hz bandwidth centered on each emission frequency (0.5f0, 1.5f0, 2.5f0, 3.5f0, f0, 2f0, 3f0, 4f0, and broadband) for each treatment target. Each mouse had 5-8 treatment targets, covering either the entire tumor or the striatum in non-tumor-bearing mice. For each target, emissions recorded during the FUS+MB treatment were normalized to the corresponding average baseline emission, resulting in a fold change over baseline. Foldchanges were averaged across targets to obtain an overall treatment-level foldchange for each emission signature.

**Tissue Processing for Flow Cytometry**

Either 3 or 7 days after completing FUS+MB treatment, mice were transcardially perfused with 10 mL of cold 1X PBS. The tumor-bearing hemisphere of the brain was harvested (cerebellum and olfactory bulbs removed) and placed in cold RPMI+FBS. The tissue was minced using a surgical blade and then passed through an 18-gauge needle to mechanically homogenize it. The homogenate was then enzymatically digested in RMPI+FBS with Collagenase A (1 mg/mL; Sigma-Aldrich #10103578001) and DNase I (1 mg/mL; Sigma-Aldrich #10104159001) at 37°C for 20 min before being passed through a 70 μm strainer (Corning) and washed with RPMI+FBS. Myelin was separated from the mononuclear cells by resuspending the cell pellet in 10 mL of 40% Percoll (Sigma-Aldrich #P1644) and centrifuging for 10 min at 650 x g. The myelin and liquid were aspirated, and the remaining cell pellet was resuspended in 10 mL RPMI+FBS followed by centrifugation at 1800 RPM for 10 min. The samples were then resuspended in 400 μL of RPMI+FBS and kept on ice until staining. Spleens were harvested in cold RPMI+FBS, mechanically homogenized, and washed through a 40 μm strainer (Corning). The sample was centrifuged at 1200 RPM for 5 min. The cell pellet was resuspended in RBC lysis buffer (ThermoFisher Scientific #00-4333-57) for 2 min before neutralizing with 13 mL of RPMI+FBS. After another 5 min centrifugation, the final cell pellet was resuspended in 500 μL RPMI+FBS and kept on ice for staining.

**Immunofluorescence**

Immunofluorescence staining was conducted on the Ventana Discovery Ultra Staining Module (Ventana Co., Tucson, AZ). 4 μm tissue sections were deparaffinized using EZ Prep and subjected to heat-induced antigen retrieval for 64 minutes in Cell Conditioning 1 (pH 8.4). Following sequential blocking of nonspecific binding with Casein (12 min) and endogenous peroxidases with CM1 (8 min), a triple-staining protocol was initiated. Sections were first incubated for 60 minutes with either Abcam mCD4 (ab183685) or eBioscience CD8a (14-0808-82) at a 1:100 dilution, then detected using OmniMap anti-rabbit/rat HRP and the DISCOVERY Rhodamine 6G kit (Ventana). Subsequently, slides were incubated for 60 minutes with Dako CD3 (A0452) at 1:100 and visualized via the OmniMap anti-rabbit HRP detection system and Cy5. Finally, Cell Signaling CD31 (77699) antibody was applied at 1:100 for 60 minutes and detected with OmniMap anti-rabbit HRP detection system and FAM. All slides were counterstained with DAPI and mounted for assessment.

Stained sections were imaged with a Zeiss LSM 880 confocal microscope (Zeiss, Germany) in sequential scanning mode using DAPI and Alexa 488, Cy5, and TRITC. Tiled images were acquired across a portion of the tumor, including some naïve brain tissue at 63X magnification. Images were analyzed with Fiji/ImageJ and staining overlap was identified using the 'Colocalization' plugin. Final images were assembled in PowerPoint (Microsoft).

#### Supplemental Figures

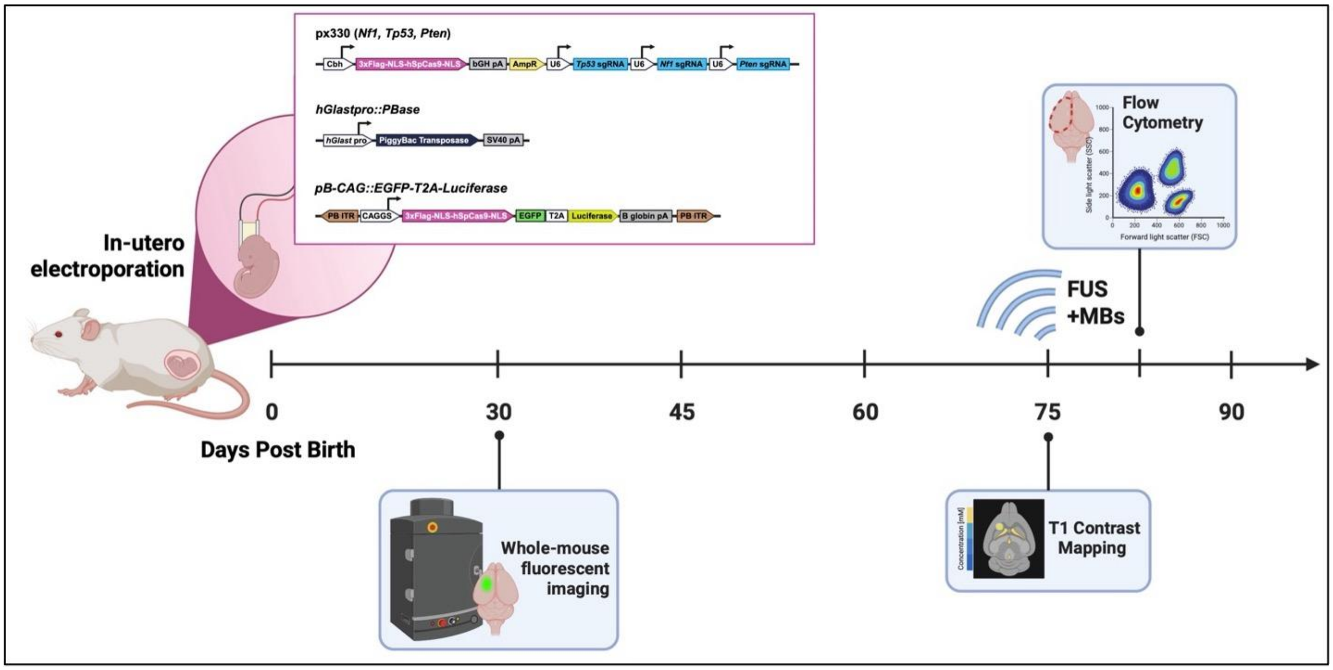


**Figure S1. 3x CRISPR GEMM Experimental Timeline.** Embryos of CD-1 IGS pregnant females underwent in utero electroporation (IUE) at embryonic day 14.5 (E14.5). Plasmids containing guide RNAs targeting the tumor suppressor genes *Nf1*, *Tp53*, and *Pten* were introduced to induce glioblastoma formation. A glial- and astrocyte-specific promoter was used to ensure targeted delivery to glial cells, and a green fluorescent protein (GFP) T2A-luciferase reporter plasmid was co-delivered to label descendant tumor cells fluorescently and allow for in vivo bioluminescence monitoring. Mice were screened at postnatal day 30 (P30) via whole-body bioluminescence imaging to confirm tumor initiation. Tumors were allowed to develop until approximately P75, when FUS+MBs was administered, followed by T1-weighted contrast mapping. Brains were harvested three days post-FUS+MB treatment, and the left hemisphere was collected for analysis of the glioma-immune landscape.

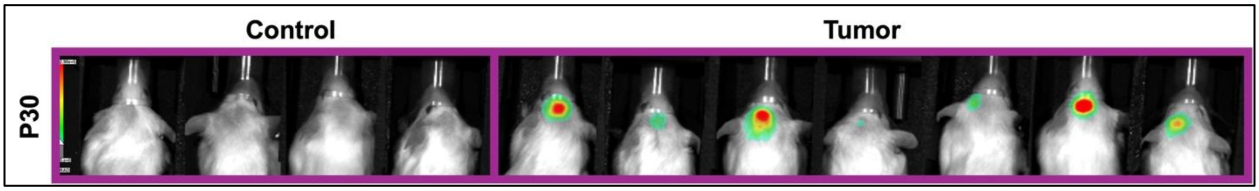


**Figure S2. Bioluminescent signal in 3x CRISPR GEMM mice from Cohort 3 on P30 for group segregation.** Luciferase activity was measured by in vivo bioluminescence imaging. The color scale 0 (no signal) to 2.90e+5 photons/sec/cm$^2$/sr (red). Control (n = 4). Tumor (n = 7).

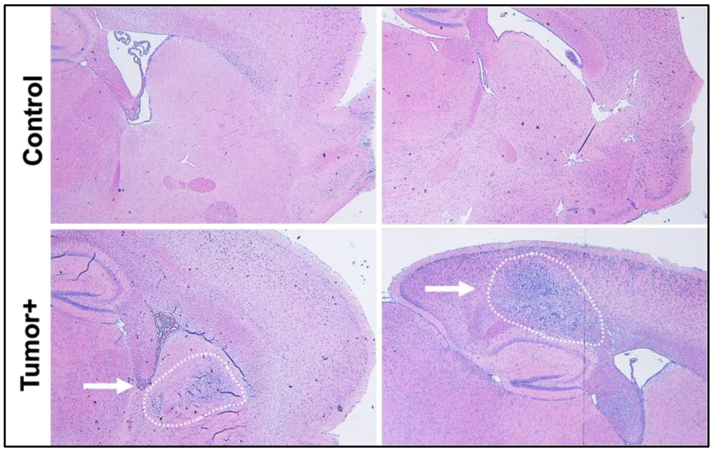


**Figure S3. Hematoxylin and Eosin (H&E) staining confirms development of 3x CRISPR tumors.** H&E-stained samples were imaged using a Zeiss microscope at 4X magnification. Representative images from Control (n = 2) and Tumor+ (n = 2) samples are shown after P65.

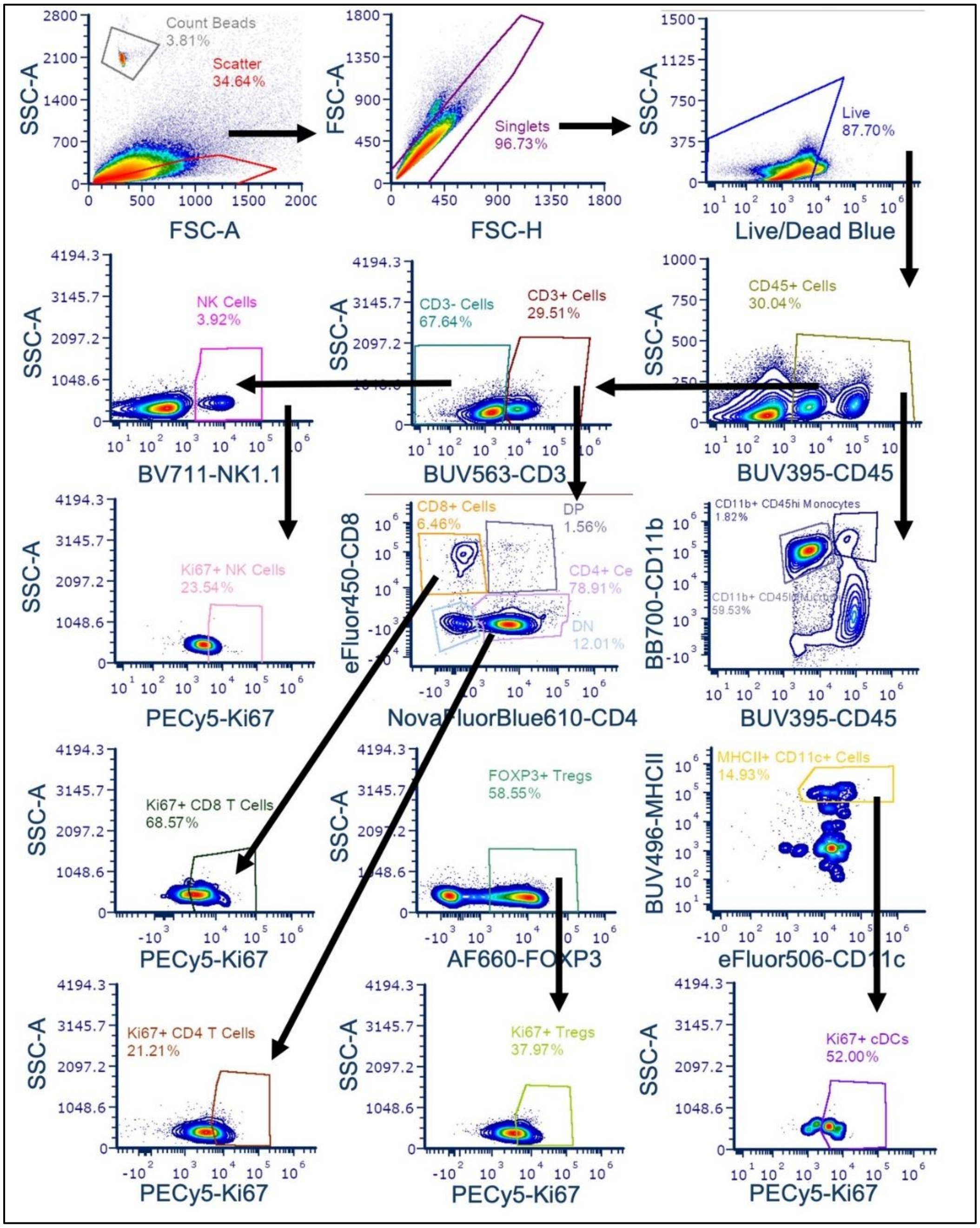


**Figure S4. Flow Cytometry gating strategy for 3x CRISPR tumor-immune landscape profiling 3 days after FUS.** Gating strategy presented for the following immune cell subsets: T cells (Live/CD45$^+$CD3$^+$); natural killer (NK) cells (Live/CD45$^+$CD3$^-$NK1.1$^+$); CD4 helper T cells (Live/CD45$^+$CD3$^+$CD4$^+$); CD8 cytotoxic T cells (Live/CD45$^+$CD3$^+$CD8$^+$); regulatory T cells (Tregs) (Live/CD45$^+$CD3$^+$CD4$^+$FOXP3$^+$); monocytes (Live/CD45$^+$CD11b$^{hi}$); dendritic cells (Live/CD45$^+$CD11b$^{hi}$CD11c$^+$MHCII$^{hi}$); microglia (Live/CD45$^+$CD11b$^{lo}$). Proliferating immune cell subpopulations were determined by Ki67 positivity. All frequencies shown are of the parent gate.

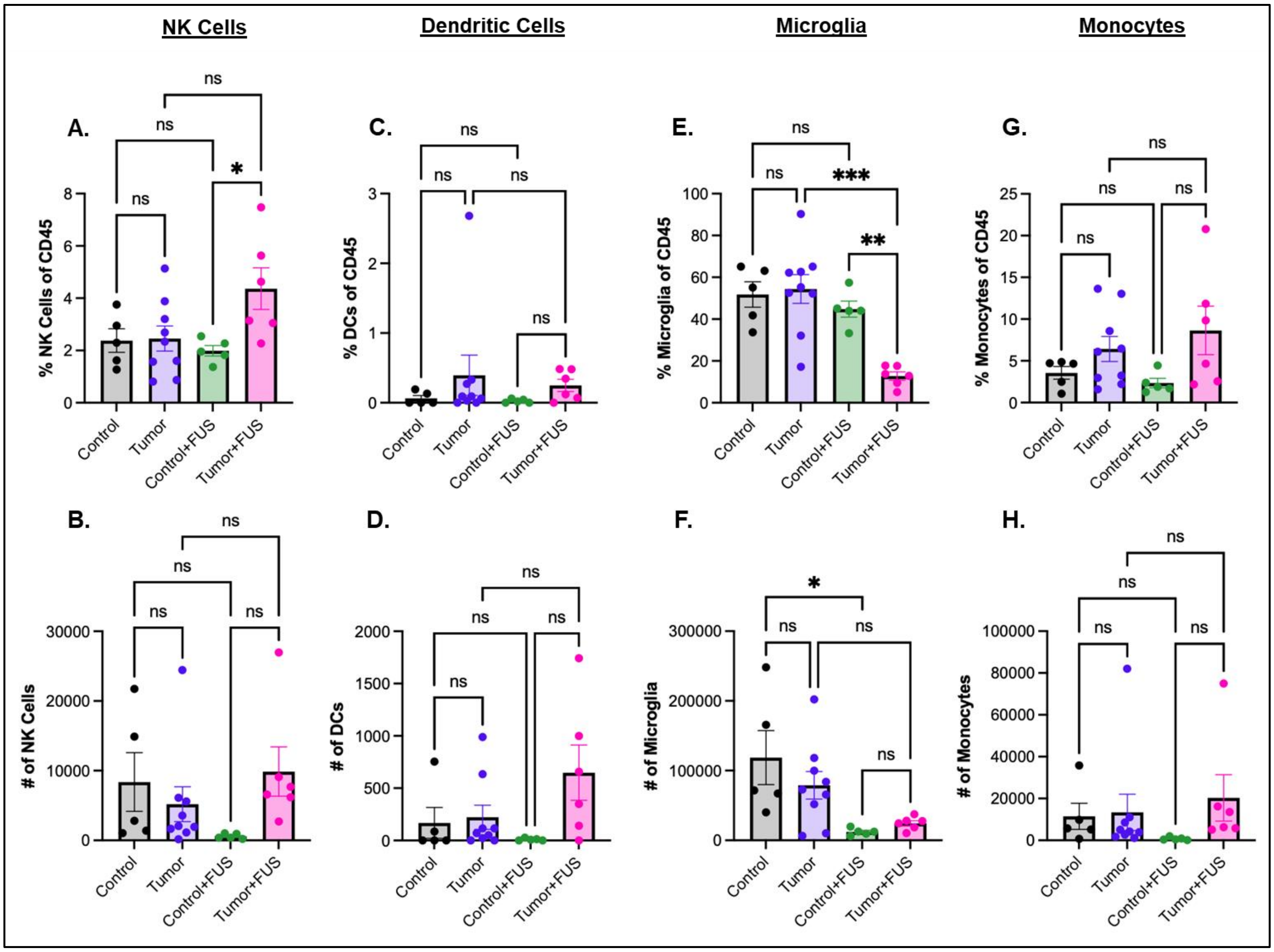


**Figure S5. FUS treatment induces minimal changes in brain NK and myeloid cell populations in 3x CRISPR mice 3 days after FUS.** A) Percent natural killer (NK) cells (Live/CD45$^+$CD3$^-$NK1.1$^+$). B) Number of NK cells. C) Percent dendritic cells (DCs) (Live/CD45$^+$CD11b$^{hi}$CD11c$^+$MHCII$^{hi}$). D) Number of DCs. E) Percent microglia (Live/CD45$^+$CD11b$^{lo}$). F) Number of microglia. G) Percent of monocytes (Live/CD45$^+$CD11b$^{hi}$). H) Number of monocytes. Two-way ANOVA with multiple comparisons correction (Tukey's). Means $\pm$ SEM. * $p < 0.05$. ** $p < 0.01$. *** $p < 0.001$. **** $p < 0.0001$.

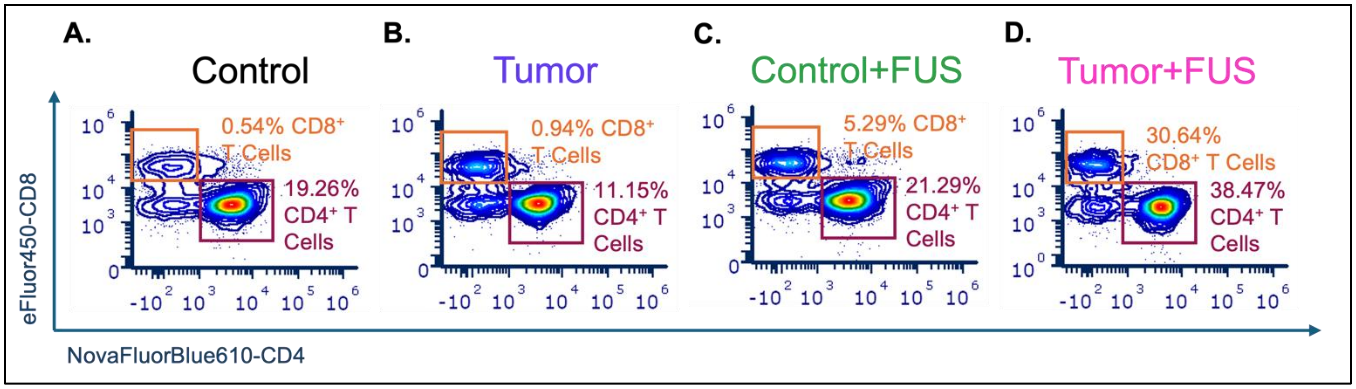


**Figure S6. Representative flow cytometry plots demonstrating enhanced T-cell infiltration in 3x CRISPR tumors following FUS treatment.** A-D) Representative flow cytometry plots showing the percentage of $CD4^+$ and $CD8^+$ T cells among $CD45^+$ immune cells in (A) control brain, (B) tumor brain, (C) control+FUS brain, and (D) tumor+FUS brain.

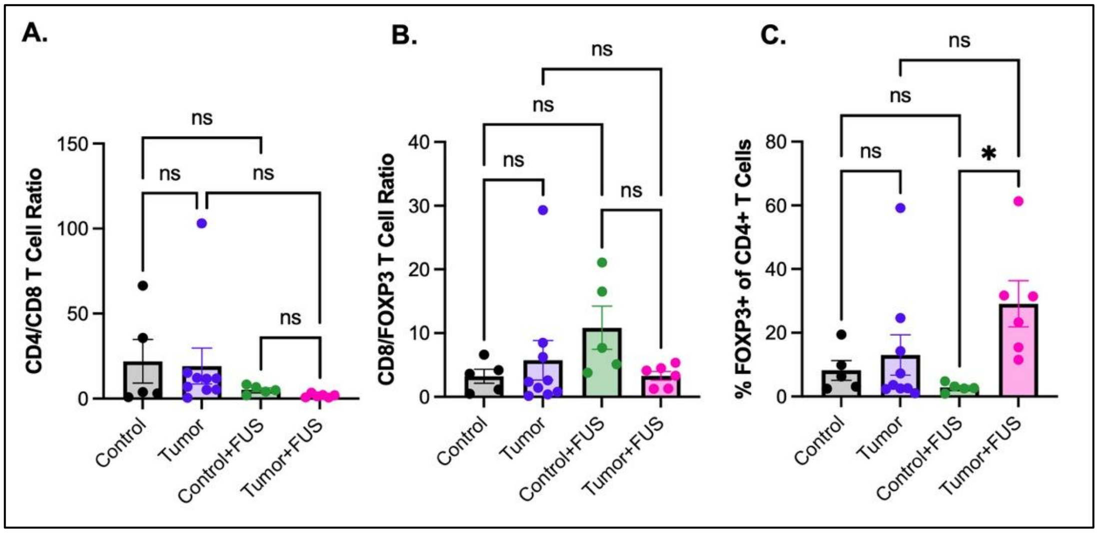


**Figure S7. FUS increases the proportion of FOXP3$^+$ regulatory T cells (Tregs) in 3x CRISPR tumors compared to control mice.** A) CD8$^+$ to CD4$^+$ T cell ratio 3 days post-FUS. B) CD8$^+$ T cell to FOXP3$^+$ regulatory T cell (Treg) ratio. C) Percent FOXP3$^+$ Tregs of CD4$^+$ T cells. Control (black, n = 5). Tumor (purple, n = 9). Control+FUS (green, n = 5). Tumor+FUS (pink, n = 6). Two-way ANOVA with multiple comparisons correction (Tukey's). Means $\pm$ SEM. * $p < 0.05$.

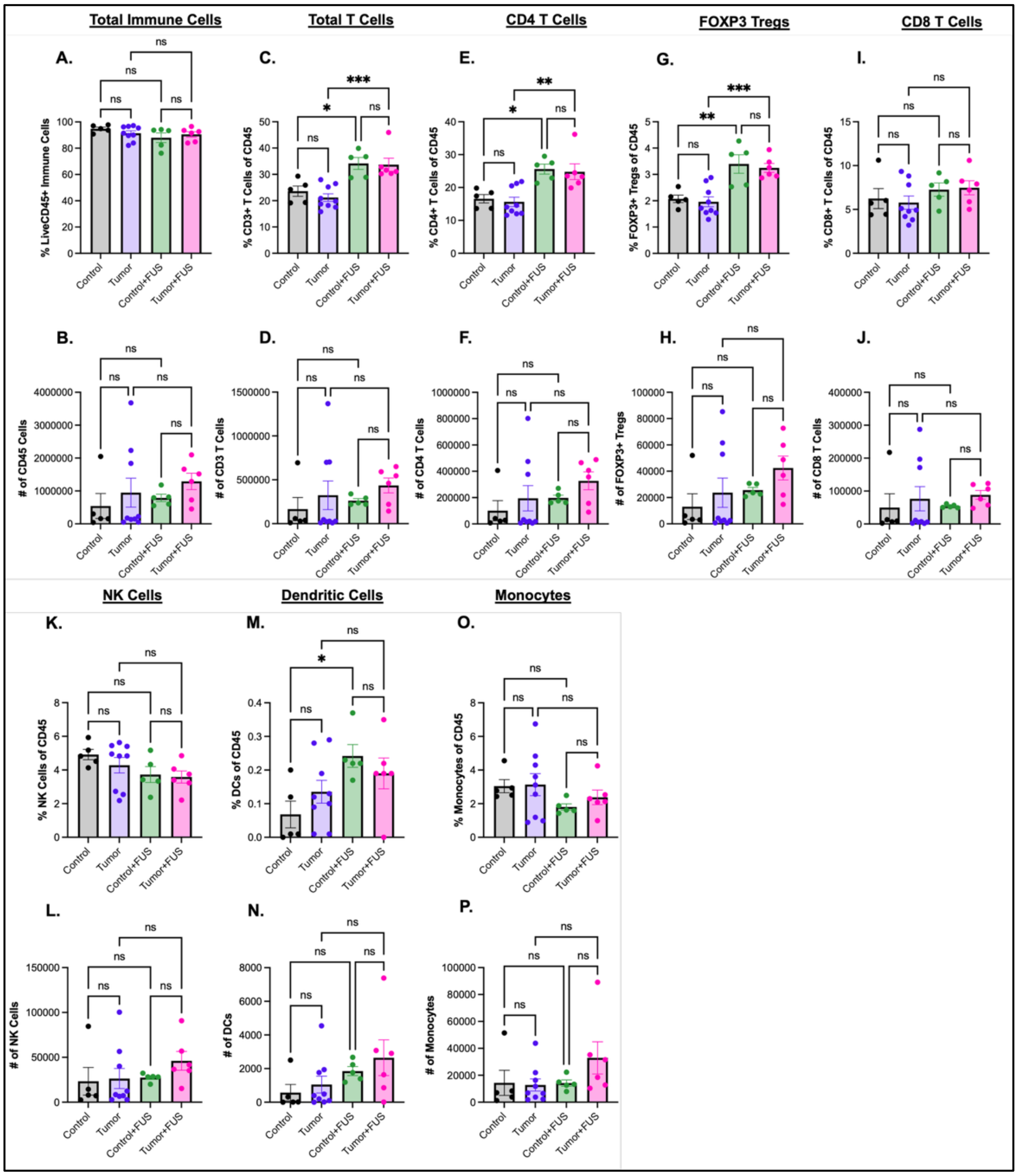


**Figure S8. Influence of FUS treatment on splenic immune cells of 3x CRISPR mice.** A) Percent total immune cells (Live/CD45$^+$). B) Number of total CD45$^+$ immune cells. C) Percent total T cells (Live/CD45$^+$CD3$^+$). D) Number of total T cells. E) Percent CD4$^+$ T cells (Live/CD45$^+$CD3$^+$CD4$^+$). F) Number of CD4$^+$ T cells. G) Percent FOXP3$^+$ regulatory T cells (Tregs) (Live/CD45$^+$CD3$^+$CD4$^+$FOXP3$^+$). H) Number of FOXP3$^+$ Tregs. I) Percent CD8$^+$ T cells (Live/CD45$^+$CD3$^+$CD8$^+$). J) Number of CD8$^+$ T cells. K) Percent double negative (DN) T cells (Live/CD45$^+$CD3$^+$CD4$^-$CD8$^-$). L) Number of DN T cells. M) Percent double positive (DP) T cells (Live/CD45$^+$CD3$^+$CD4$^+$CD8$^+$). N) Number of DP T cells. O) Percent natural killer (NK) cells (Live/CD45$^+$CD3$^-$NK1.1$^+$). P) Number of NK cells. Q) Percent dendritic cells (DCs) (Live/CD45$^+$CD11b$^{hi}$CD11c$^+$MHCII$^{hi}$). R) Number of DCs. S) Percent of monocytes (Live/CD45$^+$CD11b$^{hi}$). T) Number of monocytes. Two-way ANOVA with multiple comparisons correction (Tukey's). Means ± SEM. * $p < 0.05$. ** $p < 0.01$. *** $p < 0.001$.

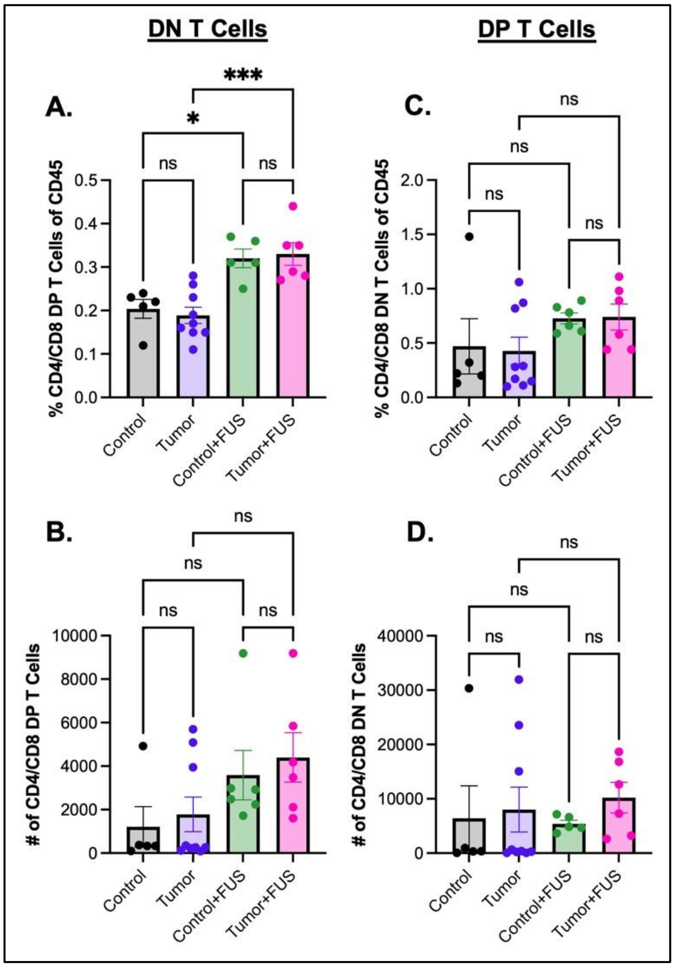


**Figure S9. Double negative (DN) and double positive (DP) T cells are present in the spleens of 3x CRISPR GEMM mice at similar frequencies.** A) Percent double negative (DN; $CD4^{-}/CD8^{-}$) T cells in the brain. B) Number of DN T cells. C) Percent double positive (DP; $CD4^{+}/CD8^{+}$) T cells in the brain. D) Number of DP T cells. Control (black, n = 5). Tumor (purple, n = 9). Control+FUS (green, n = 5). Tumor+FUS (pink, n = 6). Two-way ANOVA with multiple comparisons correction (Tukey's). Means ± SEM. * $p < 0.05$. ** $p < 0.01$. *** $p < 0.001$.

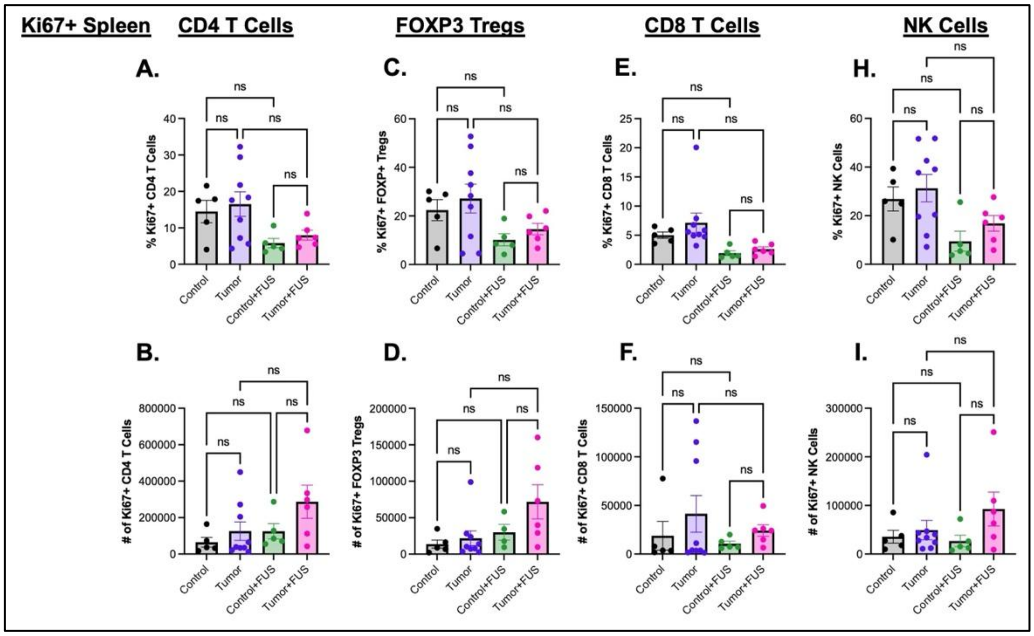


**Figure S10. 3x CRISPR mice treated with FUS show no changes in Ki67⁺ expression on immune populations in the spleen.** A) Percent Ki67⁺ CD4⁺ T cells. B) Number of CD4⁺ T cells. C) Percent Ki67⁺ FOXP3⁺ regulatory T cells (Tregs). D) Number of FOXP3⁺ Tregs. E) Percent Ki67⁺ CD8⁺ T cells. F) Number of CD8⁺ T cells. G) Percent Ki67⁺ natural killer (NK) cells. H) Number of NK cells. I) Percent Ki67⁺ dendritic cells (DCs). Two-way ANOVA with multiple comparisons correction (Tukey's). Means ± SEM. * $p < 0.05$.

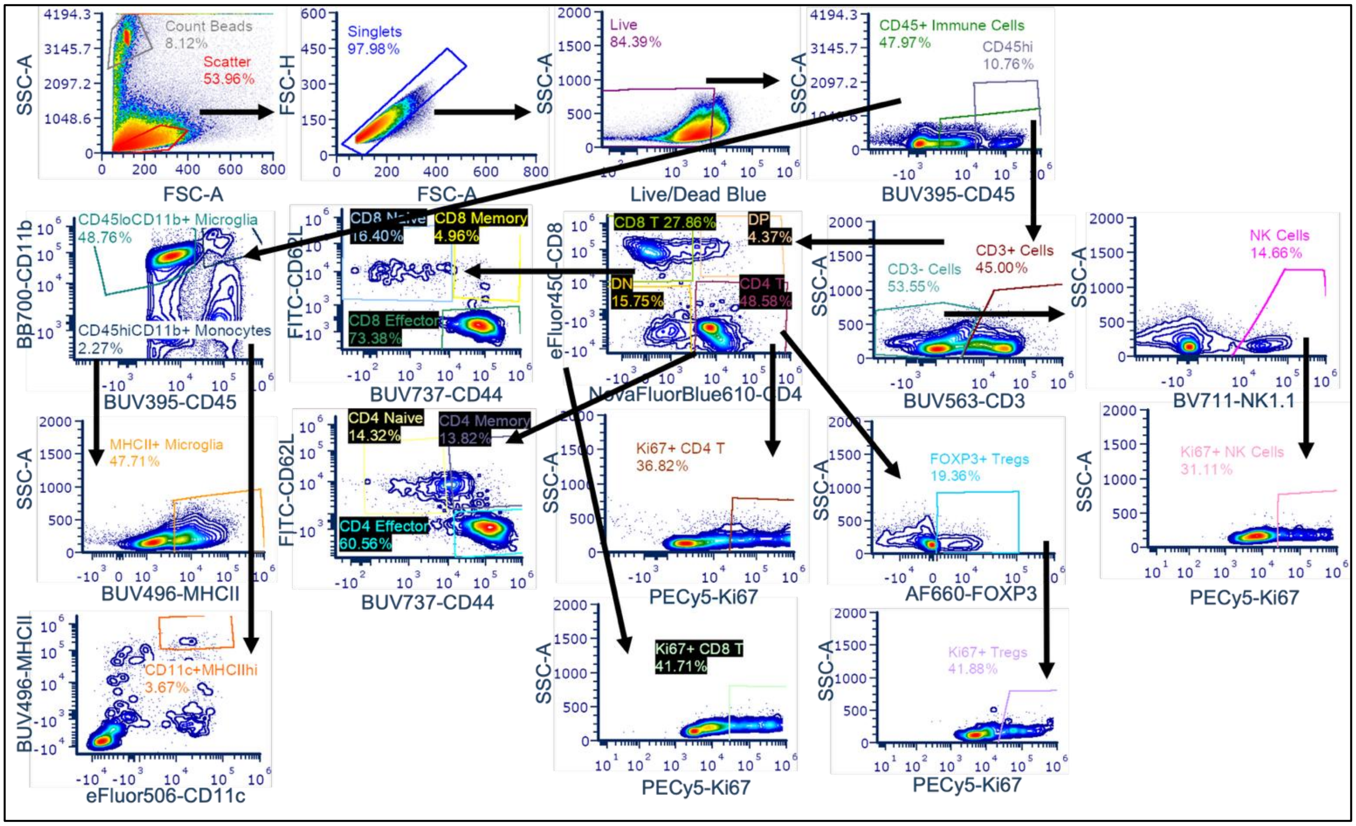


**Figure S11. Flow Cytometry gating strategy for 3x CRISPR tumor-immune landscape profiling 7 days after FUS.** Gating strategy presented for the following immune cell subsets: T cells (Live/CD45$^+$CD3$^+$); natural killer (NK) cells (Live/CD45$^+$CD3$^-$NK1.1$^+$); CD4$^+$ helper T cells (Live/CD45$^+$CD3$^+$CD4$^+$); CD8$^+$ cytotoxic T cells (Live/CD45$^+$CD3$^+$CD8$^+$); regulatory T cells (Tregs) (Live/CD45$^+$CD3$^+$CD4$^+$FOXP3$^+$); monocytes (Live/CD45$^+$CD11b$^{hi}$); dendritic cells (Live/CD45$^+$CD11b$^{hi}$CD11c$^+$MHCII$^{hi}$); microglia (Live/CD45$^+$CD11b$^{lo}$); MHCII$^+$ microglia (Live/CD45$^+$CD11b$^{lo}$MHCII$^+$); naïve T cells (Live/CD45$^+$CD3$^+$CD4/CD8$^+$CD44$^{lo}$CD62L$^{hi}$); memory T cells (Live/CD45$^+$CD3$^+$CD4/CD8$^+$CD44$^{hi}$CD62L$^{hi}$); effector T cells (Live/CD45$^+$CD3$^+$CD4/CD8$^+$CD44$^{hi}$CD62L$^{lo}$). Proliferating immune cell subpopulations were determined by Ki67 positivity. All frequencies shown are of the parent gate.

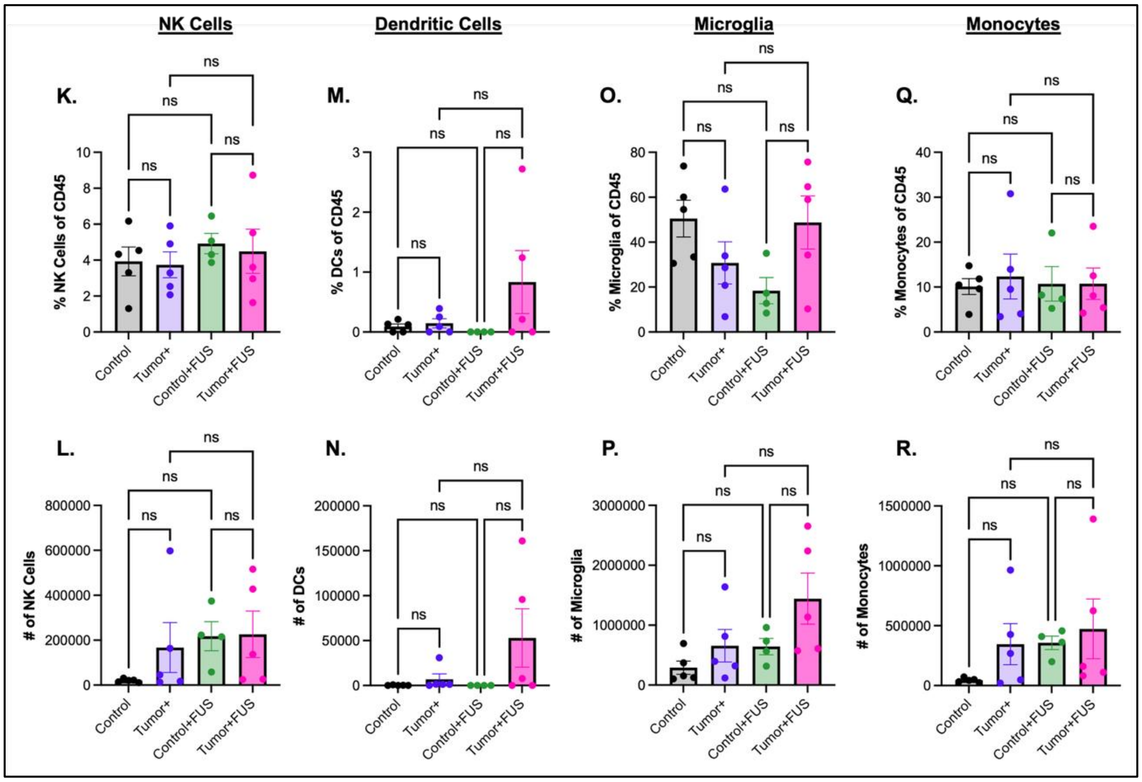


**Figure S12. Brain NK and myeloid cell populations remain unchanged 7 days after FUS treatment in 3x CRISPR mice.** A) Percent natural killer (NK) cells (Live/CD45$^+$CD3$^-$NK1.1$^+$). B) Number of NK cells. C) Percent dendritic cells (DCs) (Live/CD45$^+$CD11b$^{hi}$CD11c$^+$MHCII$^{hi}$). D) Number of DCs. E) Percent microglia (Live/CD45$^+$CD11b$^{lo}$). F) Number of microglia. G) Percent of monocytes (Live/CD45$^+$CD11b$^{hi}$). H) Number of monocytes. Two-way ANOVA with multiple comparisons correction (Tukey's). Means $\pm$ SEM.

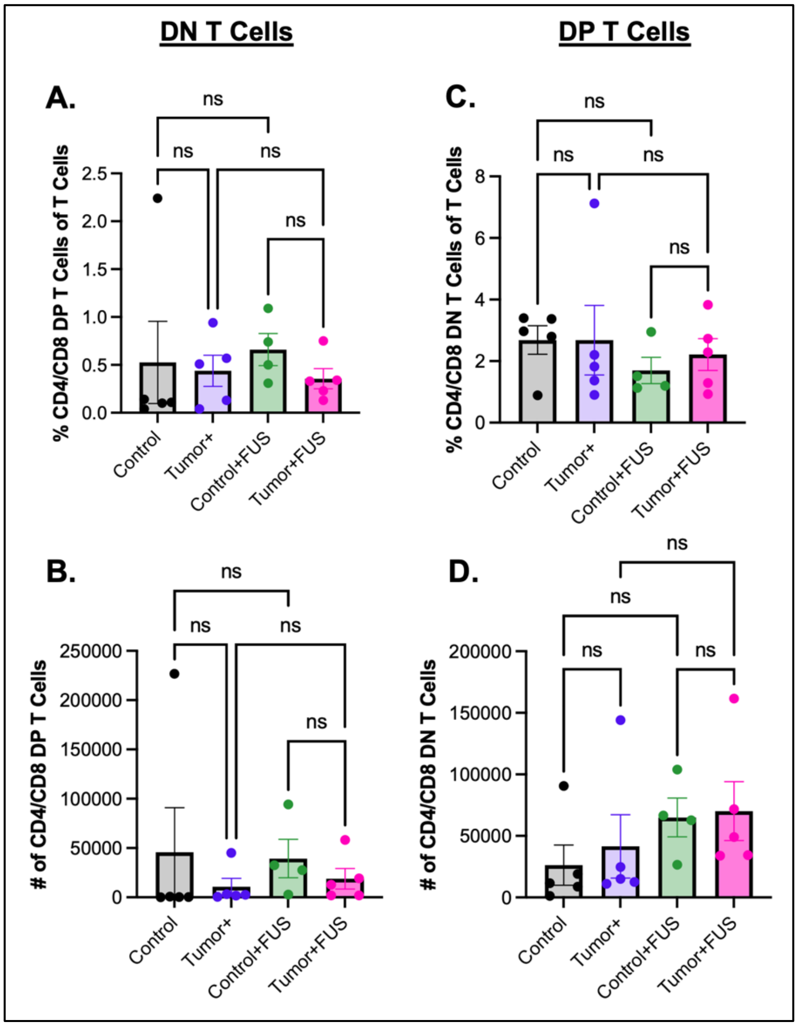


**Figure S13. Double negative (DN) and double positive (DP) T-cell populations remain comparable in 3x CRISPR GEMM tumors 7 days after FUS.** A) Percent double negative (DN; $CD4^{-}/CD8^{-}$) T cells in the brain. B) Number of DN T cells. C) Percent double positive (DP; $CD4^{+}/CD8^{+}$) T cells in the brain. D) Number of DP T cells. One-way ANOVA with multiple comparisons correction (Sidak Model). Means $\pm$ SEM.

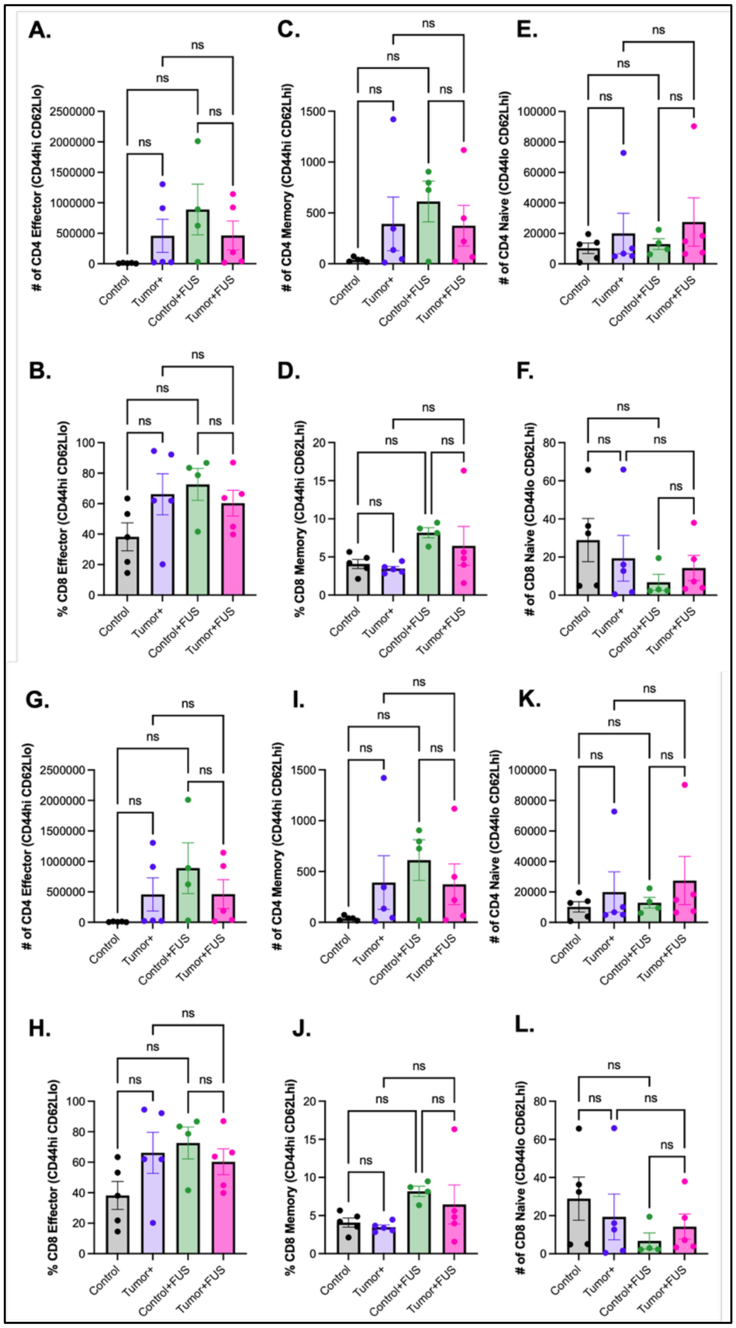


**Figure S14. FUS does not alter brain effector, naïve, or memory T-cell populations 7 days post-FUS treatment.** A) Percent CD4$^+$ Effector T cells (CD44$^{hi}$CD62L$^{lo}$). B) Number of CD4$^+$ Effector T cells (CD44$^{hi}$CD62L$^{lo}$). C) Percent CD4$^+$ Memory T cells (CD44$^{hi}$CD62L$^{hi}$). D) Number of CD4$^+$ Memory T cells (CD44$^{hi}$CD62L$^{hi}$). E) Percent CD4$^+$ Naive T cells (CD44$^{lo}$CD62L$^{hi}$). F) Number of CD4$^+$ Naive T cells (CD44$^{lo}$CD62L$^{hi}$). G) Percent CD8$^+$ Effector T cells (CD44$^{hi}$CD62L$^{lo}$). H) Number of CD8$^+$ Effector T cells (CD44$^{hi}$CD62L$^{lo}$). I) Percent CD8$^+$ Memory T cells (CD44$^{hi}$CD62L$^{hi}$). J) Number of CD8$^+$ Memory T cells (CD44$^{hi}$CD62L$^{hi}$). K) Percent CD8$^+$ Naive T cells (CD44$^{lo}$CD62L$^{hi}$). L) Number of CD8$^+$ Naive T cells (CD44$^{lo}$CD62L$^{hi}$). Two-way ANOVA with multiple comparisons correction (Tukey's). Means $\pm$ SEM.

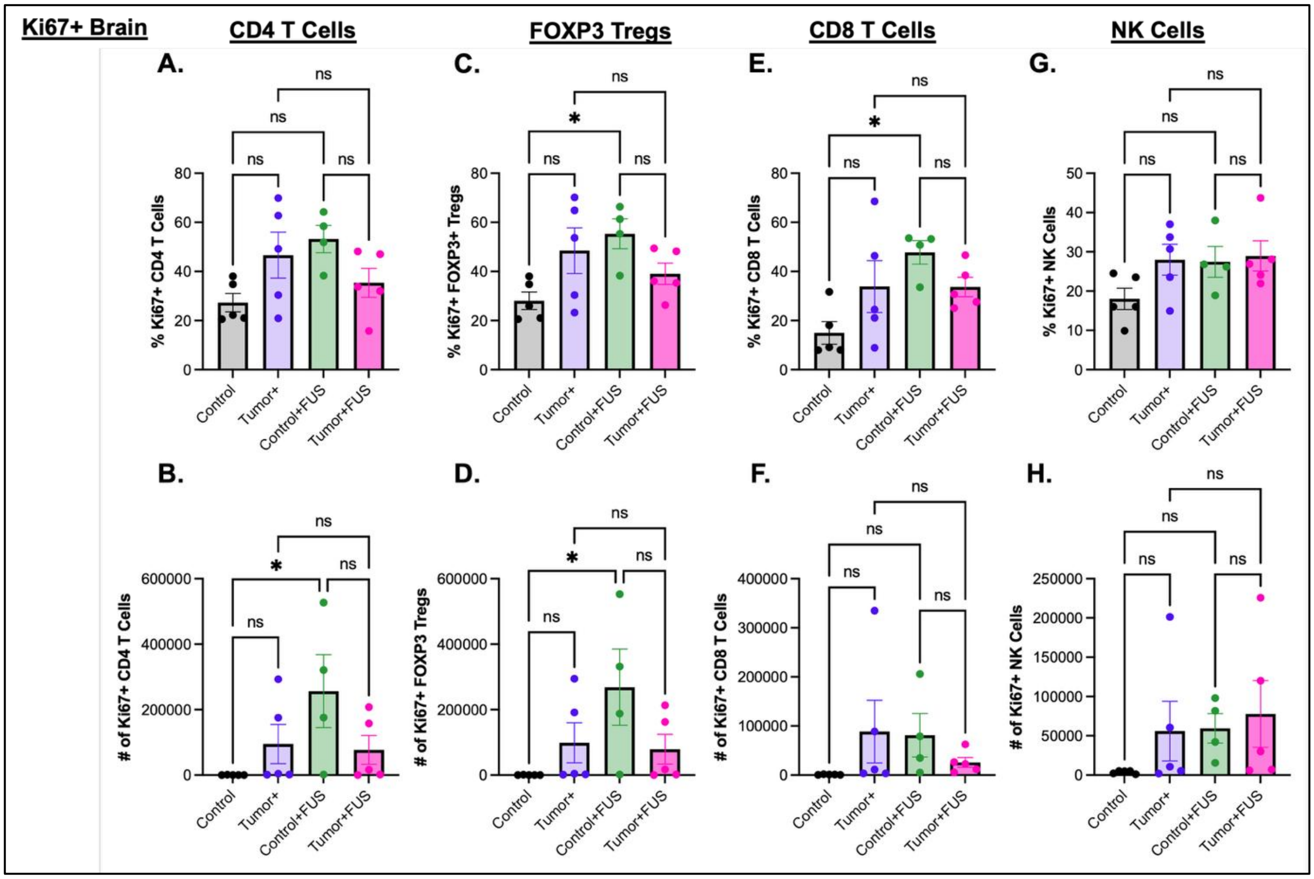


**Figure S15. Proliferation of brain immune cell populations remains largely unchanged 7 days after FUS treatment in 3x CRISPR mice.** A) Percent Ki67$^+$ CD4$^+$ T cells. B) Number CD4$^+$ T cells. C) Percent Ki67$^+$ FOXP3$^+$ regulatory T cells (Tregs). D) Number of FOXP3$^+$ Tregs. E) Percent Ki67$^+$ CD8$^+$ T cells. F) Number of CD8$^+$ T cells. G) Percent Ki67$^+$ natural killer (NK) cells. H) Number of NK cells. Two-way ANOVA with multiple comparisons correction (Tukey's). Means $\pm$ SEM.

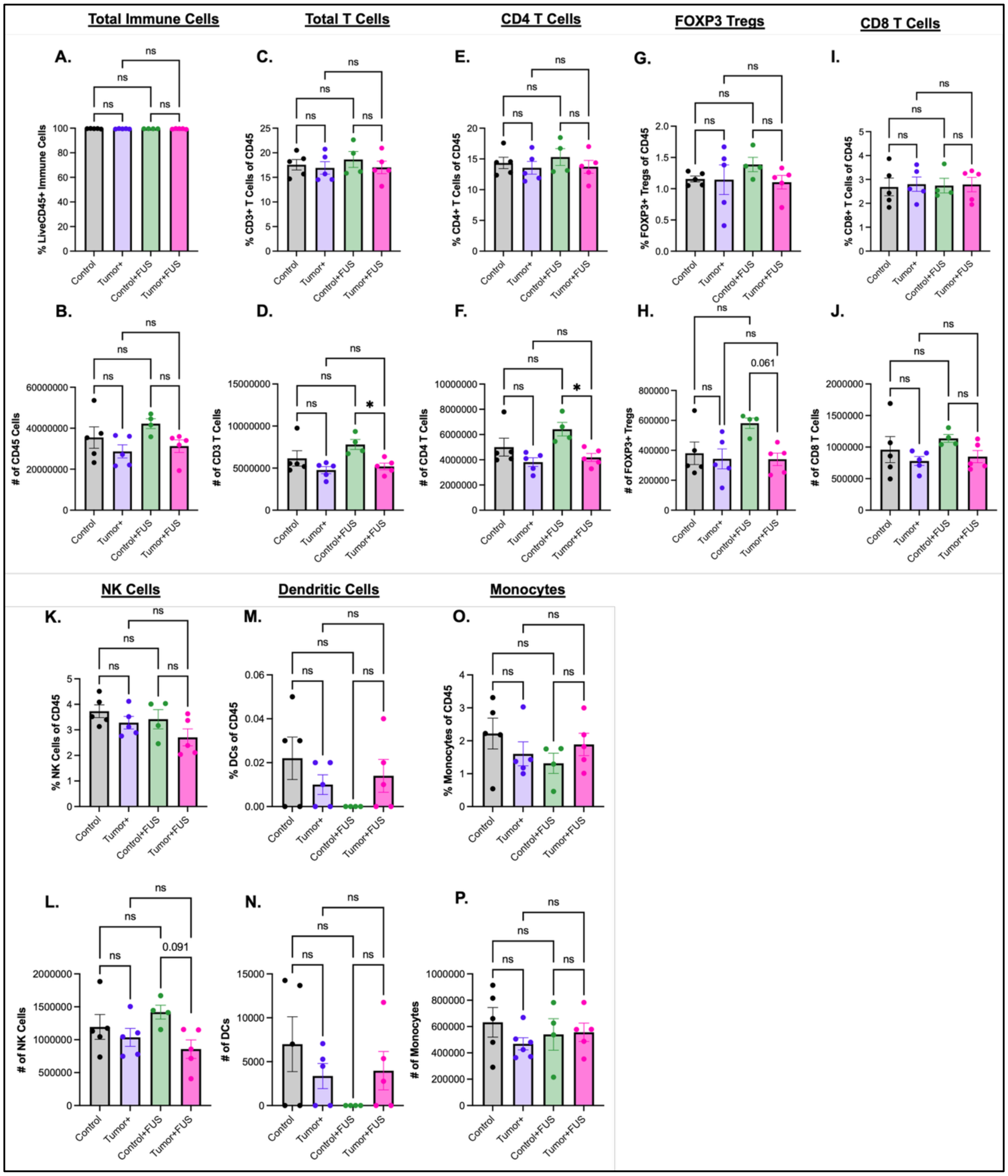


**Figure S16. Splenic immune cell populations remain largely unchanged 7 days after FUS treatment in 3x CRISPR mice.** A) Percent total immune cells (Live/CD45$^+$). B) Number of total CD45$^+$ immune cells. C) Percent total T cells (Live/CD45$^+$CD3$^+$). D) Number of total T cells. E) Percent CD4$^+$ T cells (Live/CD45$^+$CD3$^+$CD4$^+$). F) Number of CD4$^+$ T cells. G) Percent FOXP3$^+$ regulatory T cells (Tregs) (Live/CD45$^+$CD3$^+$CD4$^+$FOXP3$^+$). H) Number of FOXP3$^+$ Tregs. I) Percent CD8$^+$ T cells (Live/CD45$^+$CD3$^+$CD8$^+$). J) Number of CD8$^+$ T cells. K) Percent natural killer (NK) cells (Live/CD45$^+$CD3$^-$ NK1.1$^+$). L) Number of NK cells. M) Percent dendritic cells (DCs) (Live/CD45$^+$CD11b$^{hi}$CD11c$^+$MHCII$^{hi}$). N) Number of DCs. O) Percent of monocytes (Live/CD45$^+$CD11b$^{hi}$). P) Number of monocytes. Two-way ANOVA with multiple comparisons correction (Tukey's). Means $\pm$ SEM.

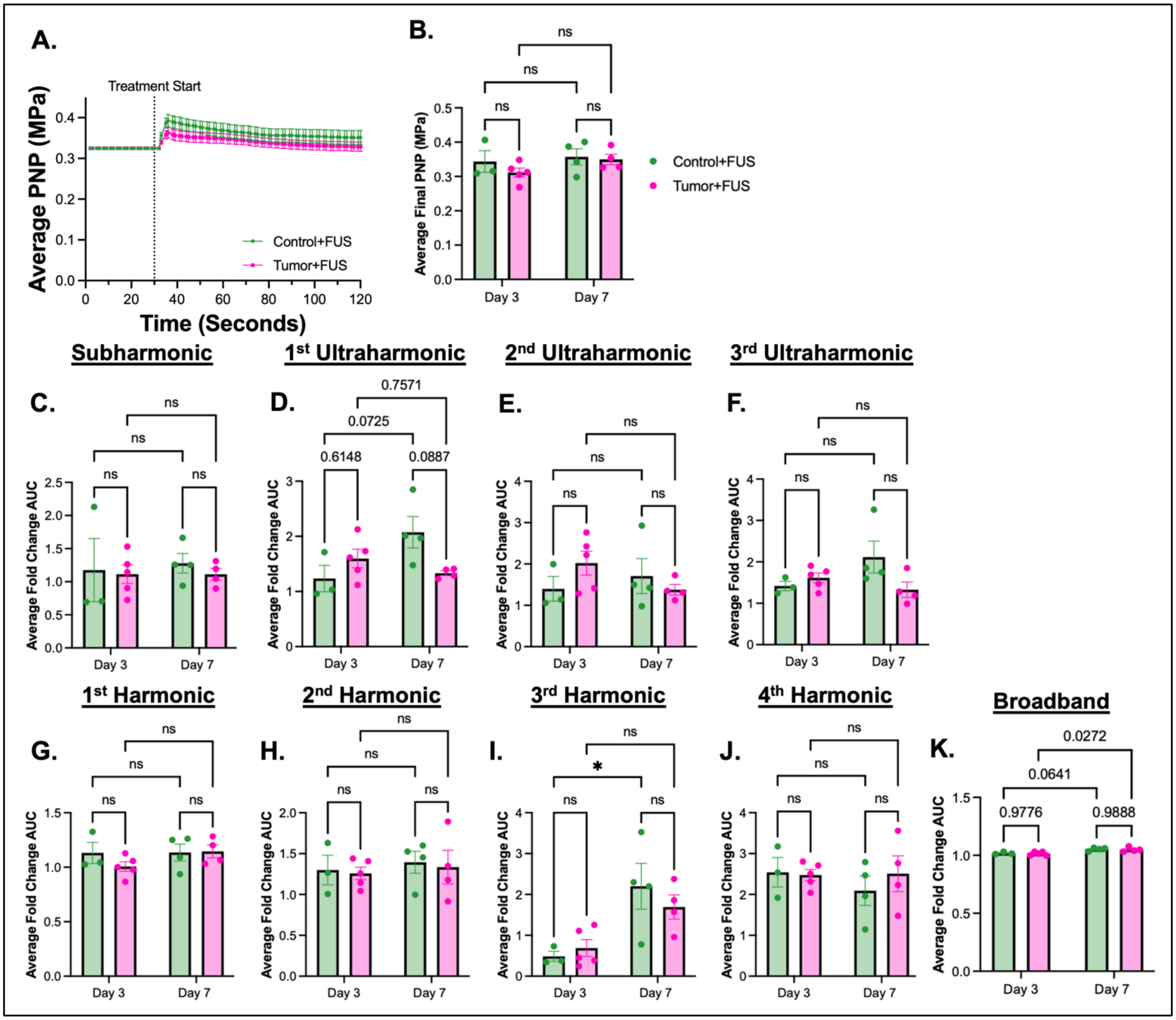


**Figure S17. PCD emissions show evidence of BTB opening in 3x CRISPR mice.** A) Average PNP (MPa) over 2-minute treatment time. B) Corresponding average final PNP. Quantification of the foldchange over baseline. C) Quantification of the foldchange over baseline emission for subharmonic. D) Quantification of the foldchange over baseline emission for 1st ultraharmonic. E) Quantification of the foldchange over baseline emission for 2nd ultraharmonic. F) Quantification of the foldchange over baseline emission for 3rd ultraharmonic. G) Quantification of the foldchange over baseline emission for 1st harmonic. H) Quantification of the foldchange over baseline emission for 2nd harmonic. I) Quantification of the foldchange over baseline emission for 3rd harmonic. J) Quantification of the foldchange over baseline emission for 4th harmonic. K) Quantification of the foldchange over baseline emission for broadband. One-way ANOVA with multiple comparisons correction (Sidak Model). Means ± SEM. * $p < 0.05$.

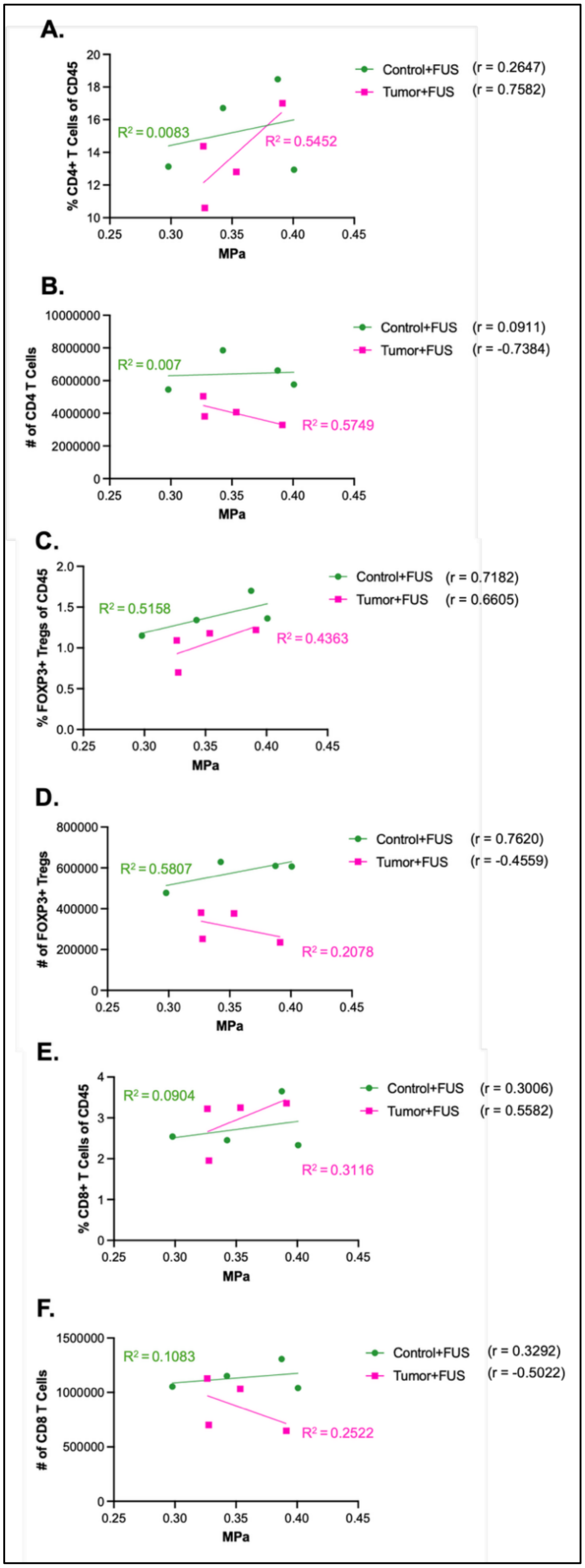


**Figure S18. Correlation between average final peak negative pressure (PNP) from FUS and T-cell responses in the 3x CRISPR mice.** A) Percent CD4 T cells vs. PNP (MPa). B) Number CD4 T cells vs. PNP (MPa). C) Percent FOXP3$^{+}$ Tregs vs. PNP (MPa). D) Number FOXP3$^{+}$ Tregs vs. PNP (MPa). E) Percent CD8 T cells vs. PNP (MPa). F) Number CD8 T cells vs. PNP (MPa). n = 4 mice per group. Correlations were assessed using Pearson's correlation coefficient (r) and $R^2$. All correlations were not statistically significant ($p > 0.05$).
